# Tailoring the self-blinking of sulfonamide rhodamine for long-term protein-localizing super-resolution imaging

**DOI:** 10.1101/2024.06.12.598600

**Authors:** Xue Zhang, Ying Zheng, Lujia Yang, Zhiwei Ye, Yi Xiao

## Abstract

Life continually changes its protein arrangements, yet the molecular ultradetails are covered by the short-lived deficiency of fluorophore blinking for super-resolution imaging. Herein, we proposed a crowding strategy to conserve the self-blinking events for prolonging the imaging time. We engineered sulfonamide rhodamines through atom-radii expansion (O-C-Si), rationally reversing xanthene intersection and creating stacking to enhance ring-opening energetical barriers. Our stacked rhodamines demonstrated decreased recruiting rates and extended survival lifetimes at single-molecule level, validating the decreased self-blinking kinetics from stacking strategy. Accordingly, our silicon-substituted rhodamine enabled persistent molecular localization imaging of various sub-organelle proteins to state-of-art time (0.5 h) in living cells, with versatile capabilities for three-dimensional and dual-color imaging. We envision our crowding strategy sets a new stage for prolongating super-resolution imaging through structural engineering.

## 1. Introduction

The secret of life embeds in the changes of biomolecule arrangement beyond diffraction limit. Super-resolution imaging, specifically the single-molecule localizing approach, paves the way to uncover the protein localizations with the aid of protein-tag technologies.^1–4^ However, the dynamics of protein arrangements confronts an imaging time barrier due to the bleaching of fluorophore blinking.^5–7^ Although exchangeable labeling was developed to bypass the fluorophore bleaching, this strategy requires homogeneously distributed fluorophores beyond the irregular naturality of living cells. The fundamental key to break the barrier is the engineering of blinking kinetics through tailoring fluorophore molecular structures. By exceedingly decreasing blinking frequency, a single molecule extends its surviving time, and these fluorophores group to prolongate imaging time and satisfy the dynamic arrangement demands. The challenge is how to rationally tune the fundamental molecular structures to conquer long-term challenge and fill the gap on molecular structure-blinking kinetic relationship.

The previous distinct approach, exchangeable labeling, lacks reliability for living-cell long-term localization imaging. Developed strategies include point accumulation for imaging in nanoscale topography (PAINT),^8–12^ high-density environmentally sensitive (HIDE) strategies^13–16^ and exchangeable fusion protein labeling.^17,18^ The core idea of exchangeable labeling is the dynamic binding between fluorophores and targets, through utilizations of the fluorescence environmental sensitivity in cases of HIDE or PAINT strategies, and the mutation of HaloTag protein like [xHTL]HaloTag7.^17^ However, exchangeable labeling encounters degressive reliability in the cell center, as the concentration of fluorophores reduces with internalization in context of cellular enclosure. Thus, it is a necessity to return to the foundation of blinking events i.e., the molecule structure to tune the blinking kinetics for long-term super-resolution imaging.

In this regard, self-blinking is an ideal selection for obtaining long-term super-resolution imaging, as self-blinkable rhodamines exhibited a thermo-dynamic dark-bright equilibrium modulable by molecular structures (Figure 1a). In a landmark work, Urano et al. firstly generalized the idea of self-blinking and developed hydroxymethyl rhodamine (HMSiR) for SMLM imaging.^19^ Later, researchers including us have focused on the energetical,^20–29^ kinetic^30^ and emissive wavelength^31–34^ context of self-blinking rhodamines. Notably, Toomre et.al utilized the environmental sensitivity of HMSiR to achieve long-term exchangeable labeling of the lipophilic membranes.^15^ Yet, all rhodamines failed to tune the blinking kinetics of self-blinking from the structural perspective, excluding the long-term imaging capability from a single fluorophore molecule. It is particularly crucial to rationalize the self-blinking rhodamine structure to modulate its blinking kinetics for achieving long-term SMLM imaging.

**Figure 1.**
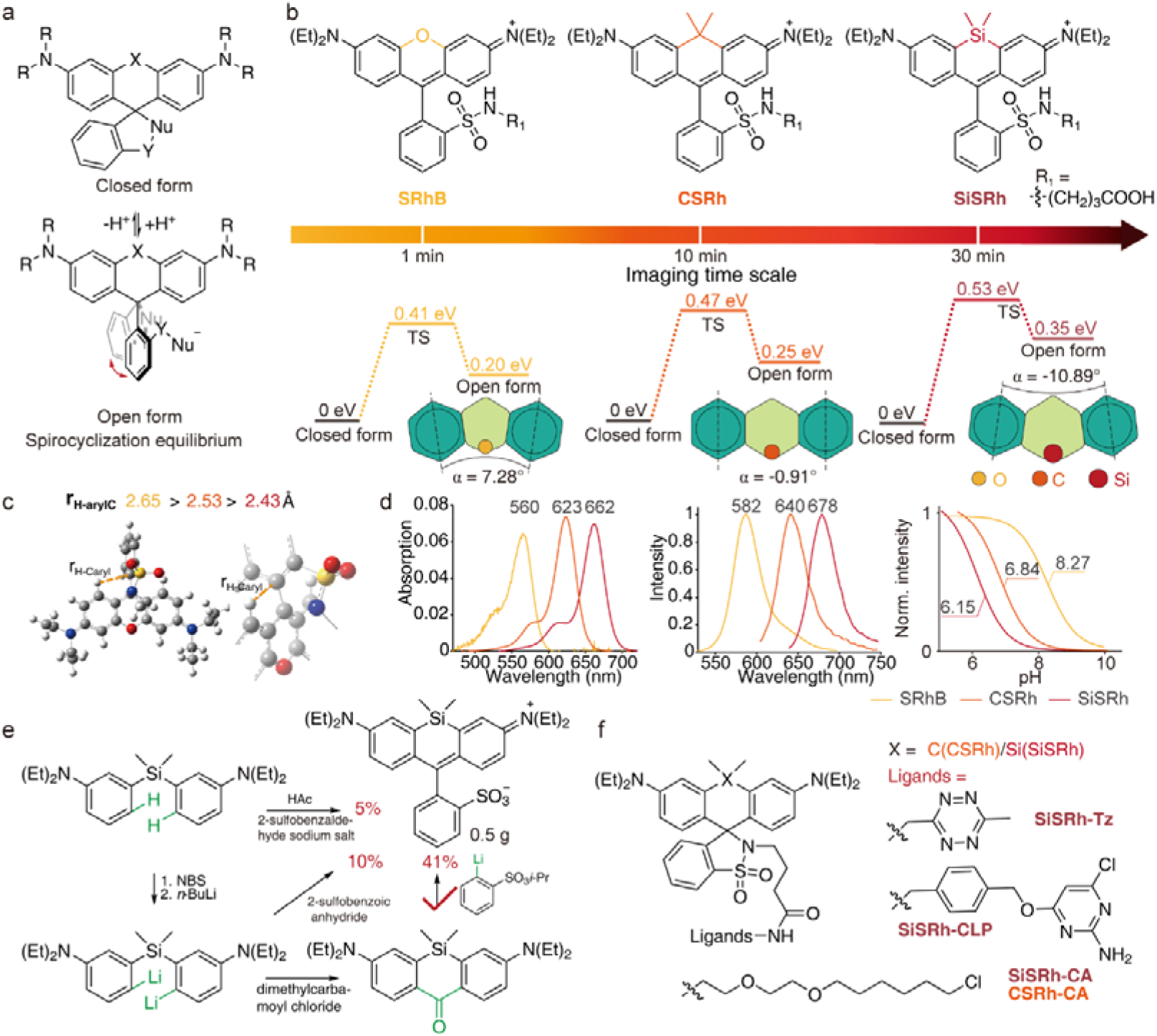
Chemistry of crowding strategy. (a) Structural transforms of the rhodamine spirocyclization equilibrium between a closed lecuo form and a ring-opened zwitterion. (b) Molecular structures of designed sulfonamide rhodamines with heteroatom-expansion (O-C-Si) with calculated ring-opening energies. Inset presented the illustrative geometry of molecules at transient states (TS). (c) Crowding induced steric effect for transient state revealed by reduced H-C_aryl_ distance. (d) Spectrum in enthanol and integrated emission intensity vs pH in PBS buffer (contained 30% enthanol). (e) Key synthetic route for Si-sulfonamide rhodamine scaffold. (f) Developed sulfonamide rhodamine probes for protein labeling.

In this study, we rationalized the steric effect of spirocyclization equilibrium of sulfonamide rhodamines. Through replacement of the xanthene heteroatom, the intersection of symmetric phenyl units was reversed, creating a crowding induced stacking and an energetic barrier to delay the ring-opening kinetics. The followingly developed sulfonamide rhodamine (SRhB), carbon sulfonamide rhodamine (CSRh) and silicon sulfonamide rhodamine (SiSRh) demonstrated decreased recruiting rates and extended survival time at single molecule level, corresponding to enhanced steric effect from their reversed crowding. More importantly, the decreased blinking kinetics leaded to significantly enlarged imaging time window for living-cell protein-based super-resolution imaging. In addition, SiSRh demonstrated state-of-art 30 minutes time window as well as multi-dimensional and dual-color imaging functionalities.

## 2. Results and Discussion

### 2.1 Probe Design

Recently, we have discovered the blinking kinetics (like recruiting rate, i.e. the ring-opening rate) of self-blinkable rhodamines determined their imaging time window instead of p*K*_cycling_ value (the negative logarithm of thermal constant of spirocyclization equilibrium).^30^ This identification of kinetic over thermal propensity driven us to rethink of the building graph of self-blinking fluorophore structure, echoing the significance of steric effect on kinetics. Thus, the problem appears that is it practicable to utilize the steric effect to control the self-blinking kinetics for extending the imaging window?

To this regard, we initially investigate the mechanics behind self-blinking. A self-blinking event corresponds to a ring-opening transform on spirocyclized rhodamine through hybridization change of C on 9-xanthene. The tetrahedral sp^3^ to planer sp^2^ transform induce the phenyl unit torsion, which could be hindered by steric effects like the crowding of nearby atoms. Thus, it is hypothesized that by increasing the atom size of the xanthene heteroatom (O 1.71 < C 1.9 < Si 2.32, Figure 1b),^35^ it is possible to expand the heterocyclic ring, create stack on the spirocyclic side, slow the spirocyclic kinetics and finally prolongate the imaging time window. To this regard, we optimized the zwitterionic and lactone structure of sulfonamide rhodamines (SRhB, CSRh, SiSRh) through computational chemistry. The result (Figure 1b, illustrative scheme) validated the reversed intersection (α = 7.28° to -10.89°) from the symmetric central axis of xanthene side phenyl units, in synchronous with the atom-radii increment, implying an enhanced stacking on the spirocyclization location. The energetic barrier of ring-opening transforms was monotonically enhanced (0.41 eV, SRhB; 0.47 eV, CSRh; 0.53 eV, SiSRh; Figure 1b) in accordance with the heteroatom sizes, inferring decreased kinetics on the ring-opening. Although the electron negativity of O-C-Si atom (3.44 > 2.55 > 1.90) might contribute to the energetic barrier, the reverse introduced a steric hindrance at the transient state (TS) from crowding between the aromatic hydrogen and the torsional phenyl units (*r*_H-arylC_ = 2.65 > 2.53 > 2.43 Å, Figure 1c). Overall, by introducing size-expanded heteroatoms for xanthene, three sulfonamide rhodamines were rationally designed with increasing ring-opening energetic barrier, expected to grant delayed self-blinking kinetics for expanding imaging time window.

Thus, we decided to synthesize the series sulfonamide rhodamines. Oxygen-bridged SRhB was conveniently obtained through acid condensation following our previous report.^2330^ However, in obtaining SiSRh, we discovered that the traditional acid-catalyzed condensation resulted in poor yields (5%, Figure 1e and S1-S2), as the high temperature resulted in large amounts of byproducts. To increase the yield, we introduced organolithium chemistry by substituting the ortho-hydrogen by bromo and later exchanging to lithium. The acquired dilithiated compound was reacted directly to 2-sulfobenzoic anhydride, yet this route still leaded to low yields of 1-10% due to the low solubility of anhydride in tetrahydrofuran. The low yields forced us to further separate the construction of the xanthene and the phenyl sulfonic acid, the dilithiated compound was reacted with dimethylcarbamoyl chloride to form the silicon-bridged xanthone, and *i*-propylsulfonylphenyl lithium was added to the xanthone to obtain the scaffold with over 40% high yields. Our results indicated the complex route might sometimes exist a superior approach than the simple ones as in the synthetic of rhodamine scaffold. By utilizing this strategy, we obtained gram level SiSRh and CSRh. At last, our obtained scaffolds of sulfonamide rhodamines were grafted protein labeling functionality through engineering Halo/SNAP-tag ligands or tetrazines (Figure 1f).

### 2.2 Spectroscopic Properties

Next, spectral study was investigated for the sulfonamide rhodamines. The absorption and emission spectra of CSRh and SiSRh are red-shifted by 60 and 100 nm respectively compared to SRhB (λ_ab_/λ_em_ = 560/582 *vs* 623/640 *vs* 662/678; order in SRhB, CSRh, SiSRh, the same follows; unit is nm, in ethanol contained 0.1% TFA, Figure 1d) with minimized crosstalk (6%) for multi-color imaging. All sulfonamide rhodamines exhibited high brightness (ε × Φ = 2.7 *vs* 4.3 *vs* 3.3; unit is × 10^4^ L·mol^-1^·cm^-1^), sufficient for single molecule imaging. p*K*_cycling_ demonstrates a gradual decrement in the order of SRhB, CSRh and SiSRh (8.27 *vs* 6.85 *vs* 6.15; Figure 1d and S3-S5), matching the theoretical ring-opening energy barriers. Accordingly, CSRh and SiSRh present decreased the spirocyclization equilibrium rates (*k*_E-a_ = 0.05 s^-1^ *vs* 0.03 s^-1^, Figure S6) for both free and fluorophore protein conjugates compared to SRhB. The spectral results demonstrate the robust brightness of sulfonamide rhodamines for multi-color localization imaging and validated the energetic barrier for ring-opening through atom-size expansion.

### 2.3 Single-Molecule Analysis

Encouraged by spectral results, the single-molecule blinking kinetics were further investigated for sulfonamide rhodamines. The typical fluorescence trajectories obtained are shown in Figure 2a. CSRh and SiSRh exhibit large number of self-blinking events following SRhB, but they demonstrate different blinking kinetics. The signals of CSRh and SiSRh molecules continuously appear during whole imaging time (presenting no signs of bleaching), whereas SRhB molecules rapidly bleach within 10 seconds. To understand this discrepancy, key single-molecule blinking kinetics are statistically summarized in Figure 2b and Table S1. Since the persistence of CSRh and SiSRh, they showed significantly prolonged last event time than SRhB (5.6 ± 0.6 *vs* 47 ± 2.2 *vs* 43 ± 4.7; unit is s, order in SRhB, CSRh, SiSRh and the same follows), suggesting many CSRh and SiRh molecules survived to the imaging end. The average dwell times of the dark-state (dark time) demonstrate an increasing order for the three (0.4 ± 0.1 vs 2.4 ± 0.4 vs 5.2 ± 0.2; unit is s, Figure 2c), and the duty cycle (the ratio of bright state dwell time versus the whole imaging time; 0.014 *vs* 0.009 *vs* 0.004, Figure 2d) exhibits a decreasing order. The prolonged dark time and small duty cycle values indicate CSRh and SiRh conserve their fluorophores in dark states with decreased probability of dark-bright transform, showing a delayed blinking kinetics in consistent with the hindered ring-opening kinetics prediction from the energy barriers. Moreover, the recruiting rate (*k*_rc_) measures the absolute ring-opening speed at ground state, and exhibits sharply decreased values (3.07 ± 0.64 *vs* 0.45 ± 0.11 *vs* 0.17 ± 0.01; unit is s^-1^, Figure 2e). The reduction of *k*_rc_ substantially confirmed the heteroatom size expansion decreases the ring-opening kinetics, indicating the delay effect from crowding strategy. Furthermore, the delayed kinetics induce significantly increased half-life time *(τ*_1/2_ = 1 *vs* 16 *vs* 57; unit is s, Figure 2f) of the fraction of survival molecules, making long-term imaging possible for CSRh and SiSRh. In addition, sulfonamide rhodamines present single-molecule brightness > 120 photons/10 ms, sufficient for localization imaging. The heteroatom size expansion from SRhB, CSRh to SiSRh results in a butterfly effect on slowing the blinking kinetics, and among the three, SiSRh exhibit mostly delayed recruiting rates with full potential for long-term super-resolution imaging.

**Figure 2.**
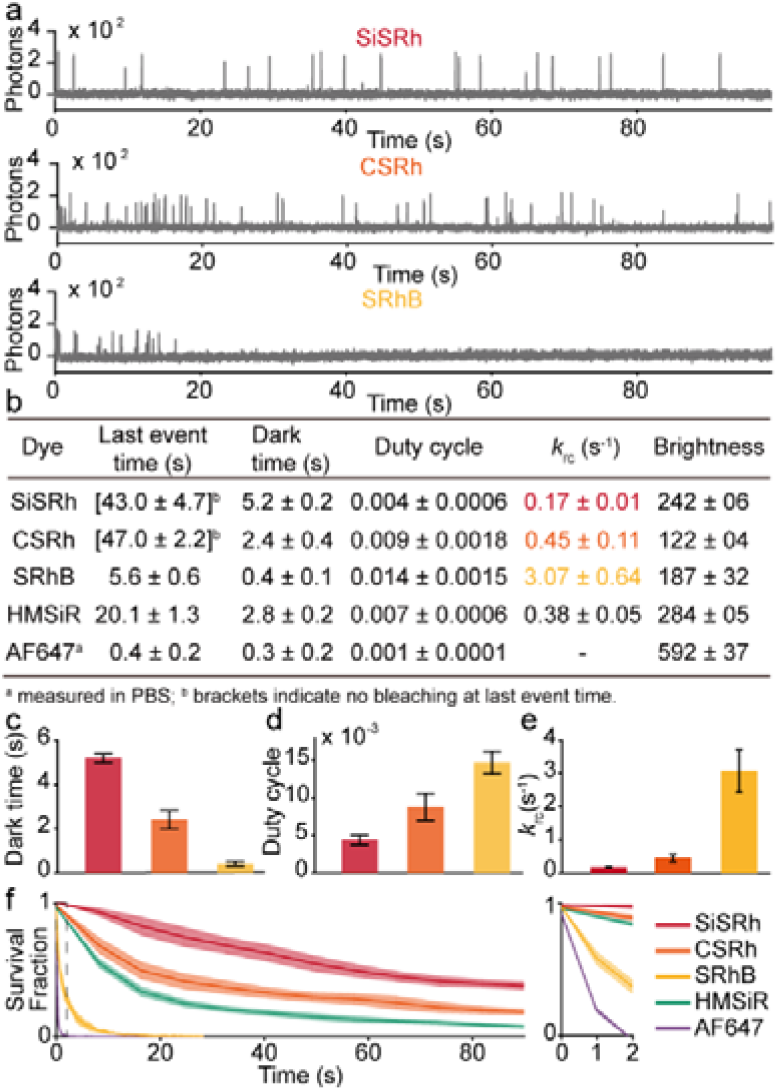
Single-molecule photophysical studies of sulfonamide rhodamines. (a) Typical fluorescence trajectories. (b) Summarization of the single-molecule characteristics between the three sulfonamide rhodamines and two cutting-edge fluorophores, HMSiR and AF647. Comparation of dark time (c), duty cycle (d) and recruiting rate (*k*_rc_, e) of the three. (f) Time evolvement changes of molecular survival fraction for investigated fluorophores. The irradiation power of all studies was 800 W/cm^2^.

To further exemplify the delayed blinking kinetics, two cutting-edge dyes AF647 and HMSiR were compared in the single-molecule study. The self-blinking propensity of SiSRh and HMSiR results in a similar large number of switching events (9.2 ± 1.1, SiSRh *vs* 8.0 ± 0.9, HMSiR; Figure 2b) in physiological buffer (phosphate buffered saline, PBS). Since SiSRh exhibits slower recruiting rate than that of HMSiR (0.38 ± 0.05 s^-1^), the former demonstrates a longer lifetime (43 ± 4.7, SiSRh vs 20.1 ± 1.3, HMSiR; unit is s) highly potential for expanding the imaging time window. In sharp contrast, AF647 was easily bleached and presented barely blink events (0.4 ± 0.2, bleach time; 1.8 ± 0.1, switch number) in PBS without the assistance of imaging enhancing. Sulfonamide rhodamines, especially SiSRh, demonstrate a high potential for long-term imaging in physiological environments.

### 2.4 Long-term living-cell super-resolution imaging

We further explored the protein-based super-resolution imaging capabilities of SiSRh, CSRh and SRhB in living cells. SiSRh, CSRh and SRhB were facilely engineered with the Halo-tag protein ligands. The constructed probes present robust membrane permeability and with the aid of Halo-tag fusion protein technology, they successful labelled proteins of interest within 2 h (Figure S7-S10). During imaging, all sulfonamide rhodamines solidly demonstrate a large number of spontaneous blinking events, and the event localization revealed the characteristic proteins arrangements (TOMM20, Sec61β, α-tubulin and H2B, Figure 3, S7-13) reflecting diversified structures of mitochondrial outer membrane, endoplasmic reticulum (ER), microtubules and nucleus. The structural protein arrangements are acquired with high localization precision (> 9.7 nm; Figure S14). Yet, a prominent persistence disparity was discovered, as SiSRh extended the imaging window of mitochondrial outer membrane and endoplasmic reticulum up to 30 min (Figure 3a and d) with satisfactory reconstructing integrity (mitochondrial fusion and endoplasmic reticulum morphology evolvement, Supplementary Movie 1, 2), whereas the same imaging time of CSRh or SRhB was reduced to 10 min (Figure 3b, e and supplementary Movie 3, 4) or even 1min (Figure 3c and f) respectively. Quantitatively, the SiSRh molecules exhibit a survival half-time of 7.3/25.6 min for mitochondrial/ER imaging, substantially superior over those of CSRh and SRhB (mitochondria, 1.8 min *vs* < 0.5 min; ER, 4.5 min *vs* < 0.5 min; order in CSRh, SRhB; Figure 3g, h). A same prolongation on imaging time window is observed in the cases of nucleus H2B (50 min, SiSRh vs 3 min, CSRh; Figure S11, S12) and α-tubulin proteins (10 min, SiSRh vs 1 min, CSRh; Figure S13). Sulfonamide rhodamines unambiguously demonstrate super-resolution imaging capability for protein arrangements, with an imaging time window in synchronous with their heteroatom expansion induced crowded stacking. Among the three, SiSRh exhibits an unprecedented time window for imaging protein arrangements, superior than the classical self-blinking dye (HMSiR, survival half-time < 10 s, Figure S15-16).

**Figure 3.**
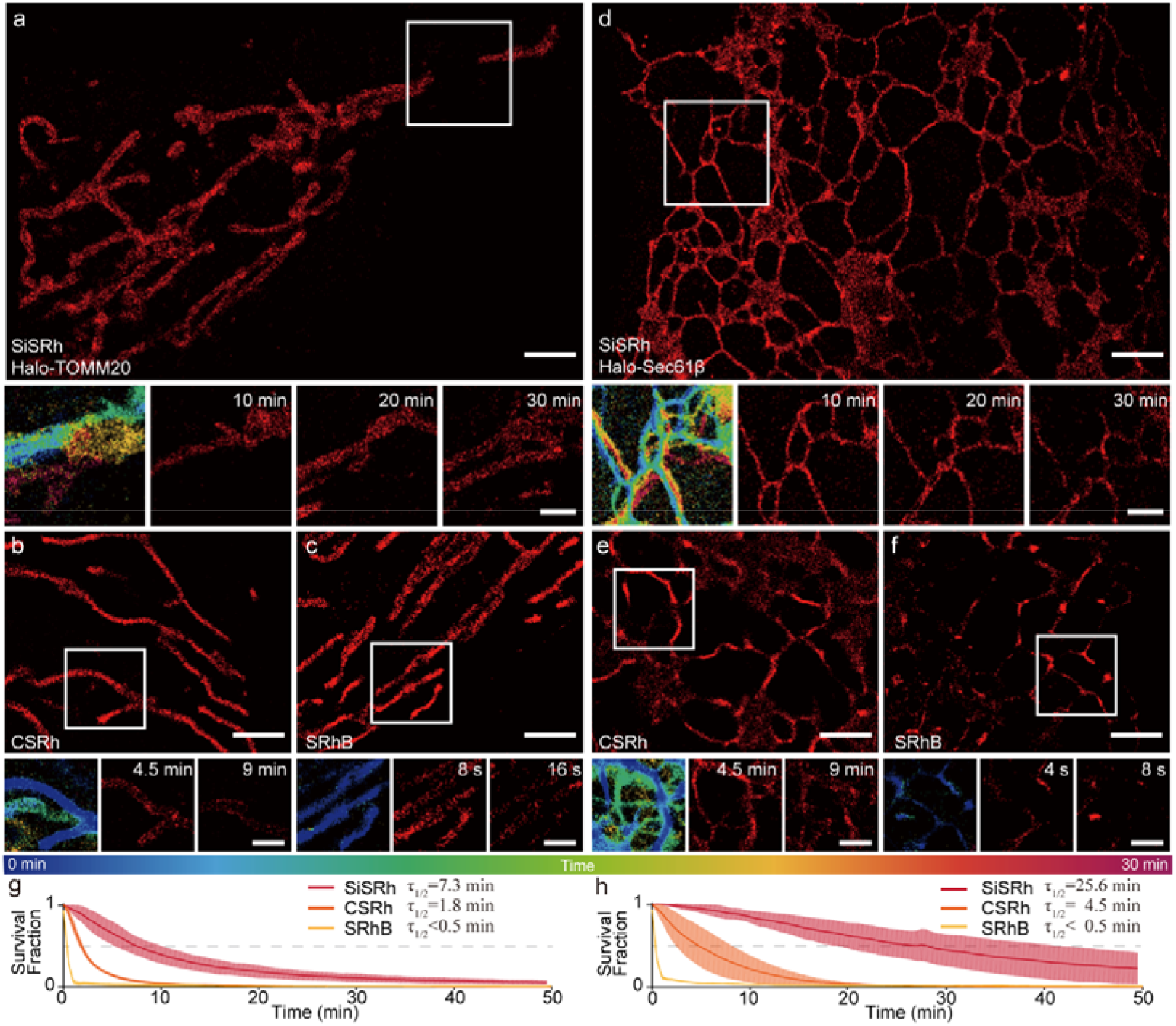
Long-term protein-localizing super-resolution imaging comparison between sulfonamide rhodamines. Reconstructions of mitochondrial TOMM20 (a-c) and endoplasmic reticulum (ER) Sec61β (d-f) in live U2OS cells through Halo-tag technology. On each panel bottom, from left to right: sequence overlay colored in 0-30 min range from three extracted reconstructions; typical reconstructions of boxed regions at indexed time point. Sulfonamide rhodamines enabled super-resolution imaging movie (0.5 h *vs* 10 min *vs* 4 s; order in SiSRh, CSRh, SRhB, the same after). The temporal resolution is 20 s, 10 s, 4 s for mitochondria and 10 s, 5s, 4 s for ER at 100 Hz acquisition. Comparison of the time evolvement of molecular survival fraction for imaging mitochondria (g) and ER (h). Scale bars: 2 μm (a, b, c, d, e, f). 1 μm (insets).

Encouraged by the Halo-tag protein results, we were interest of the universality of our crowding strategy. Thus, versatile labeling strategies (SNAP-tag, tetrazine-cyclooctyne biorthogonal strategy and immunostaining) were investigated for SiSRh in long-term super-resolution imaging. Regardless of labeling strategy, SiSRh effectively displays persistent single-molecule blinking events, enabling the reconstruction of mitochondrial morphological changes in living cells (Figure S17-20). The long-term imaging by SNAP-tag technology indicates the universal decreased blinking kinetics of SiSRh despite the protein surface environment. The application in small-molecule labeling by tetrazine-cyclooctyne biorthogonal strategy reveals the adaptivity of this fluorophore (Figure S19, S20). In fixed-cell immunolabeling study, SiSRh persistently provided blinking events over 1 h in simple PBS buffer, whereas the blinking events of commercial standard AF647 bleached in 2 min (Supplementary movie S5). The reconstruction of SiSRh accurately reproduced the filamentous structure of microtubules (localization precision: 13.4 nm) comparable to that of AF647 in imaging buffer (Figure S21-23). The durability of SiSRh during fixed-cell immunostaining further validates the general decreased blinking kinetics of SiSRh.

Thus, the crowding strategy constructed SiSRh exhibits a universality for long-term super-resolution imaging regardless of the labeling approach and microenvironment.

### 2.5 Dimensional expansion of long-term living-cell imaging

The proteins arrangement is dynamic not only in x, y plane, but also in three dimensions (*x, y, z*), which requires long-term three-dimensional (3D) imaging. The lifting from two-dimensional (2D) to 3D demands superiorly sparse single-molecule blink signals, as the axial information is acquired from the patterns of point spread function of a single fluorophore, easily distorted by background and nearby blink signals.^36–40^ The decreased blink kinetics of SiSRh enabled sparse blink events for long-term three-dimensional imaging in live cell. Through SiSRh and Halo-tag technology, the distribution of outer membranal TOMM20 proteins on the axial pore of the mitochondrial cross section was monitored for over 21 min (Figure 4a-c). In the magnified region, an emergence of TOMM20 protein clusters was discovered at 7 min, indicating the formation of a new bud from previous mitochondria. Over time, the buds grown and retracted with the formed double parallel pores subsequently reverting to a single pore at 14 min. These dynamic protein localizations accurately captured the three-dimensional mitochondrial stretching and fusion, quantitatively reported the changes of mitochondrial axial dimensions (Figure 4c). SiSRh is qualified for first revealing the dynamic pore of mitochondrial out membrane with long-time imaging capabilities for living-cell 3D super-resolution imaging.

**Figure 4.**
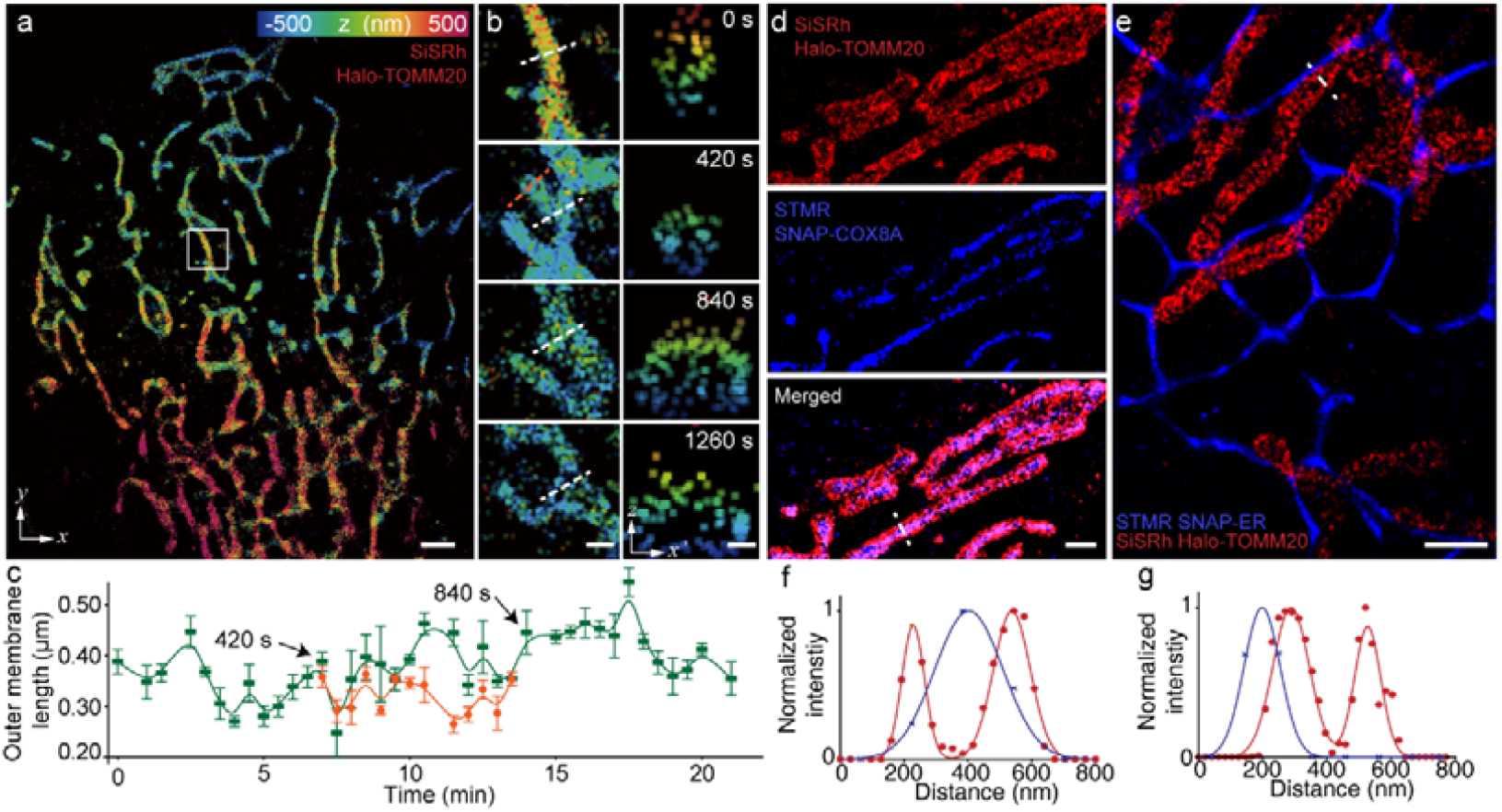
Expanding the living-cell long-term super-resolution imaging dimensions with SiSRh. The fluorophore enabled 21 min three dimensional (3D) super-resolution movie of mitochondrial outer membranes and one reconstruction (10 s, 100 Hz) at initial time was shown in (a). (b) The time-lapse magnification of boxed region in (a) with x–z projection view on the right. The dynamics of TOMM20 axial locations reveal a changing values of mitochondrial outer diameters. (d, e) Dual-color reconstruction capturing the transient interactions between mitochondrial outer membranes (TOMM20) and mitochondrial inner matrix (COX8A) or ER (specific peptides)^45^ through SiSRh and sulfonamide tetramethyl rhodamine. Intensity profiles at marked white lines, revealing the spatially separated inner and out mitochondrial membranes (f) or the significant ties between mitochondria and ER. Scale bars: 2 μm (a). 0.5/0.2 μm (b, left/right). 1 μm (d, e).

The color constitutes another dimension for acquiring interactions between biomolecules. The challenge for two-color localization imaging is the color crosstalk. Sulfonamide rhodamines spanning 550-650 nm is favorable for constructing pairs for dual-color imaging.^41–44^ Thus, SiSRh and STMR (sulfonamide tetramethyl rhodamine)^30^ was selected as the pair with minimized color crosstalk (blue channel: 565-605 nm; red channel: 667.5-725.5 nm). The reconstruction from this system successfully separates the inner outer context of mitochondria (SiSRh-Halo-TOMM20 and STMR-SNAP-COX8A) and the entanglement between mitochondria and ER (STMR-SNAP-ER; Figure 4d and e). The cross-sectional line profiles clearly distinguish the inner and outer membranal protein clusters on the mitochondria (Figure 4f), and indicate the entangled mitochondrial outer membrane on the endoplasmic reticulum (Figure 4g). Moreover, under two-color conditions, SiSRh exhibits long-term image for 10 min and resolve structural features (Figure S26). Benefited from the delayed blink kinetics from the crowding strategy, SiSRh is capable for long-term protein-based super-resolution imaging even with expanded dimensions.

With the aid of SiSRh, it is practicable to uncover the fate of proteins under photodynamic therapy. During treatment by DUT870 (a previous reported photosensitizer drug analogue),^46^ the mitochondrial outer membrane TOMM20 proteins arranged to clusters to form the division (Figure 5a and b). The specific locations arrayed at a ∼ 2 μm length for long mitochondria, echoing the worm mitochondrial origin.^47–49^ It is inferred that the clusters of TOMM20 mostly occurs at the specific polar sides of single mitochondrial worm unit. 3D imaging further supported that with the therapy elongation, the massive matrix formed in the plane projection was discovered to be individually assembled mitochondrial worm units on the axial projection (Figure 5c and d). Even under stressed microenvironments, SiSRh fulfills long-term imaging capabilities.

**Figure 5.**
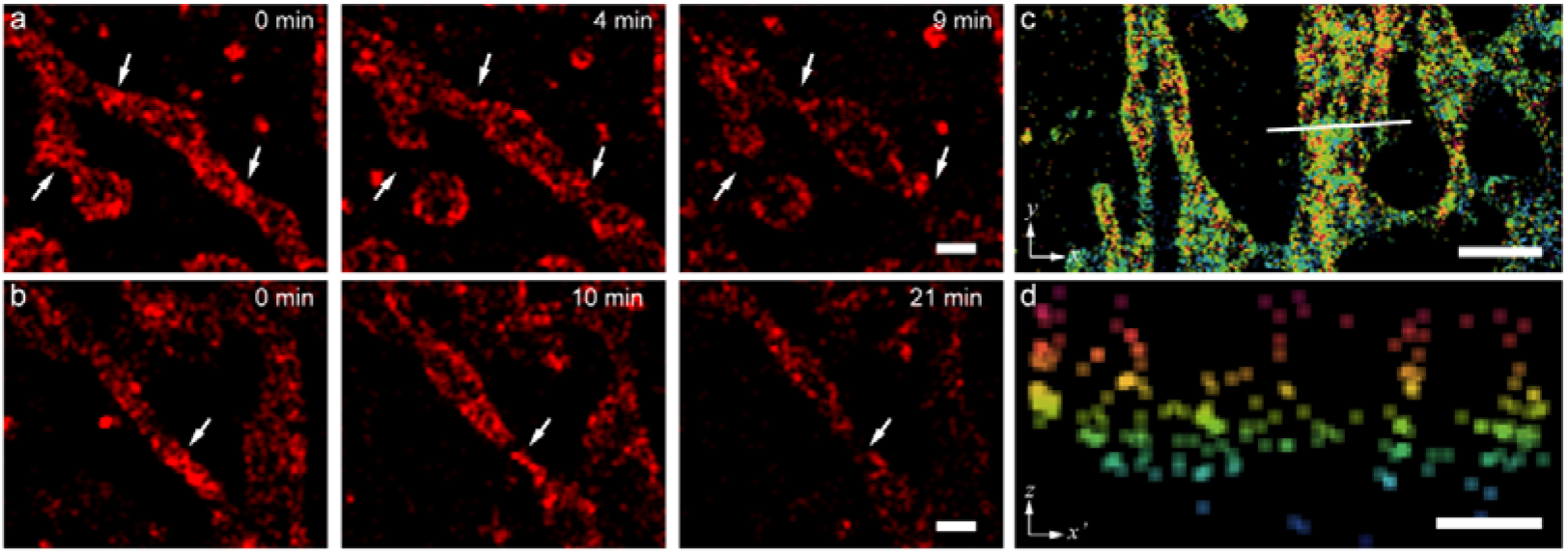
Mitochondria play worm-unit behaviors under photodynamic therapy. The super-resolution reconstructions of TOMM20 proteins through SiSRh during photodynamic therapy of a photosensitizer (DUT870). (a,b) Mitochondrial fissions at TOMM20 protein clustered locations during treatment. (c) 3D reconstructions reveal the separated worm units of mitochondrial massive matrix formed upon stressing. (d) The x–z projection view of the white line in (c). Scale bars: 2 μm (c); 0.5 μm (a, b, d).

## 3 Conclusion

In conclusion, we rationally introduced heteroatom expansion to the sulfonamide rhodamines to tailor the self-blinking kinetics. The atom-radii increment from O to C to Si resulted in the reverse of xanthene intersection (α = 7.28° to -10.89°) in a computation study, creating a steric effect by crowding stack and following an 0.12 eV enhancement on ring-opening energetical barrier. The crowding strategy decreased single-molecule blink kinetics with a > 18-fold decreased recruiting rate and 3.5-fold reduced duty cycle for SiSRh compared to SRhB. More importantly, the long-lived blinking was validated in living-cell protein-based super-resolution imaging on diversified proteins of interest, regardless of the microenvironments and labeling approaches. Beneficial from the crowding strategical delay, SiSRh exhibited a state-of-art time window (> 0.5 h) for persistently living-cell super-resolution imaging protein localizations, with versatile capability for four-dimensional and dual-color imaging. The construction of SiSRh offers a demanded probe for long-term localization imaging. We anticipate that our crowding strategy would set a new stage for developing advanced imaging agents on the perspective from steric effect and kinetics.

## Notes

### Competing Interest Statement

The authors have declared no competing interest.

## REFERENCES

(1) Xue, L.; Karpenko, I. A.; Hiblot, J.; Johnsson, K. Imaging and Manipulating Proteins in Live Cells through Covalent Labeling. Nat. Chem. Biol. 2015, 11 (12), 917–923.

(2) Minoshima, M.; Reja, S. I.; Hashimoto, R.; Iijima, K.; Kikuchi, K. Hybrid Small-Molecule/Protein Fluorescent Probes. Chem. Rev. 2023, 124 (10), 6198–6270.

(3) Reck-Peterson, S. L.; Yildiz, A.; Carter, A. P.; Gennerich, A.; Zhang, N.; Vale, R. D. Single-Molecule Analysis of Dynein Processivity and Stepping Behavior. Cell 2006, 126 (2), 335–348.

(4) Lukinavičius, G.; Umezawa, K.; Olivier, N.; Honigmann, A.; Yang, G.; Plass, T.; Mueller, V.; Reymond, L.; Corrêa, I. R.; Luo, Z. G.; Schultz, C.; Lemke, E. A.; Heppenstall, P.; Eggeling, C.; Manley, S.; Johnsson, K. A Near-Infrared Fluorophore for Live-Cell Super-Resolution Microscopy of Cellular Proteins. Nat. Chem. 2013, 5 (2), 132–139.

(5) Kwon, J.; Elgawish, M. S.; Shim, S. H. Bleaching-Resistant Super-Resolution Fluorescence Microscopy. Adv. Sci. 2022, 9, 2101817.

(6) Demchenko, A. Photobleaching of Organic Fluorophores: Quantitative Characterization, Mechanisms, Protection. Methods Appl. Fluoresc. 2020, 8, 022001.

(7) Strack, R. Bypassing Bleaching with Fluxional Fluorophores. Nat. Methods 2019, 16 (5), 357.

(8) Sharonov, A.; Hochstrasser, R. M. Wide-Field Subdiffraction Imaging by Accumulated Binding of Diffusing Probes. Proc. Natl. Acad. Sci. U. S. A. 2006, 103 (50), 18911–18916.

(9) Danylchuk, D. I.; Moon, S.; Xu, K.; Klymchenko, A. S. Switchable Solvatochromic Probes for Live-Cell Super-Resolution Imaging of Plasma Membrane Organization. Angew. Chemie - Int. Ed. 2019, 58 (42), 14920–14924.

(10) Aparin, I. O.; Yan, R.; Pelletier, R.; Choi, A. A.; Danylchuk, D. I.; Xu, K.; Klymchenko, A. S. Fluorogenic Dimers as Bright Switchable Probes for Enhanced Super-Resolution Imaging of Cell Membranes. J. Am. Chem. Soc. 2022, 144 (39), 18043–18053.

(11) Ye, Z.; Yang, W.; Zheng, Y.; Wang, S.; Zhang, X.; Yu, H.; Li, S.; Luo, C.; Peng, X.; Xiao, Y. Integrating a Far-Red Fluorescent Probe with a Microfluidic Platform for Super-Resolution Imaging of Live Erythrocyte Membrane Dynamics**. Angew. Chemie - Int. Ed. 2022, 61 (45), e202211540.

(12) Kwon, J.; Elgawish, M. S.; Shim, S. H. Bleaching-Resistant Super-Resolution Fluorescence Microscopy. Adv. Sci. 2022, 9 (9).

(13) Chu, L.; Tyson, J.; Shaw, J. E.; Rivera-Molina, F.; Koleske, A. J.; Schepartz, A.; Toomre, D. K. Two-Color Nanoscopy of Organelles for Extended Times with HIDE Probes. Nat. Commun. 2020, 11 (1), 4271.

(14) Gupta, A.; Rivera-Molina, F.; Xi, Z.; Toomre, D.; Schepartz, A. Endosome Motility Defects Revealed at Super-Resolution in Live Cells Using HIDE Probes. Nat. Chem. Biol. 2020, 16 (4), 408–414.

(15) Takakura, H.; Zhang, Y.; Erdmann, R. S.; Thompson, A. D.; Lin, Y.; McNellis, B.; Rivera-Molina, F.; Uno, S. N.; Kamiya, M.; Urano, Y.; Rothman, J. E.; Bewersdorf, J.; Schepartz, A.; Toomre, D. Long Time-Lapse Nanoscopy with Spontaneously Blinking Membrane Probes. Nat. Biotechnol. 2017, 35 (8), 773–780.

(16) Zheng, S.; Dadina, N.; Mozumdar, D.; Lesiak, L.; Martinez, K. N.; Miller, E. W.; Schepartz, A. Long-Term Super-Resolution Inner Mitochondrial Membrane Imaging with a Lipid Probe. Nat. Chem. Biol. 2024, 20 (1), 83–92.

(17) Kompa, J.; Bruins, J.; Glogger, M.; Wilhelm, J.; Frei, M. S.; Tarnawski, M.; D’Este, E.; Heilemann, M.; Hiblot, J.; Johnsson, K. Exchangeable HaloTag Ligands (XHTLs) for Multi-Modal Super-Resolution Fluorescence Microscopy. J. Am. Chem. Soc. 2023, 145 (5), 3075–3083.

(18) Holtmannspötter, M.; Wienbeuker, E.; Dellmann, T.; Watrinet, I.; Garcia-Sáez, A. J.; Johnsson, K.; Kurre, R.; Piehler, J. Reversible Live-Cell Labeling with Retro-Engineered HaloTags Enables Long-Term High- and Super-Resolution Imaging. Angew. Chemie - Int. Ed. 2023, 62 (18), e202219050.

(19) Uno, S. N.; Kamiya, M.; Yoshihara, T.; Sugawara, K.; Okabe, K.; Tarhan, M. C.; Fujita, H.; Funatsu, T.; Okada, Y.; Tobita, S.; Urano, Y. A Spontaneously Blinking Fluorophore Based on Intramolecular Spirocyclization for Live-Cell Super-Resolution Imaging. Nat. Chem. 2014, 6 (8), 681–689.

(20) Chi, W.; Tan, D.; Qiao, Q.; Xu, Z.; Liu X. Spontaneously Blinking Rhodamine Dyes for Single-Molecule Localization Microscopy. Angew. Chemie - Int. Ed. 2023, 62 (39), e2023060.

(21) Xu, Z.; Chi, W.; Qiao, Q.; Wang, C.; Zheng, J.; Zhou, W.; Xu, N.; Wu, X.; Jiang, X.; Tan, D.; Liu, X. Descriptor ΔGC[O Enables the Quantitative Design of Spontaneously Blinking Rhodamines for Live[Cell Super[Resolution Imaging. Angew. Chemie Int. Ed. 2020, 132 (45), 20390–20398.

(22) Lardon, N.; Wang, L.; Tschanz, A.; Hoess, P.; Tran, M.; D’Este, E.; Ries, J.; Johnsson, K. Systematic Tuning of Rhodamine Spirocyclization for Super-Resolution Microscopy. J. Am. Chem. Soc. 2021, 143 (36), 14592–14600.

(23) Zheng, Y.; Ye, Z.; Xiao, Y. Subtle Structural Translation Magically Modulates the Super-Resolution Imaging of Self-Blinking Rhodamines. Anal. Chem. 2023, 95 (8), 4172–4179.

(24) Martin, A.; Rivera-Fuentes, P. A General Strategy to Develop Fluorogenic Polymethine Dyes for Bioimaging. Nat. Chem. 2024, 16 (1), 28–35.

(25) Chi, W.; Qi, Q.; Lee, R.; Xu, Z.; Liu, X. A Unified Push–Pull Model for Understanding the Ring-Opening Mechanism of Rhodamine Dyes. J. Phys. Chem. C 2020, 124 (6), 3793–3801.

(26) Remmel, M.; Scheiderer, L.; Butkevich, A. N.; Bossi, M. L.; Hell, S. W. Accelerated MINFLUX Nanoscopy, through Spontaneously Fast-Blinking Fluorophores. Small 2023, 19 (12), 2206026.

(27) Remmel, M.; Scheiderer, L.; Butkevich, A. N.; Bossi, M. L.; Hell, S. W. Accelerated MINFLUX Nanoscopy, through Spontaneously Fast[Blinking Fluorophores. Small 2023, 19 (12), 2206026.

(28) Qi, Q.; Chi, W.; Li, Y.; Qiao, Q.; Chen, J.; Miao, L.; Zhang, Y.; Li, J.; Ji, W.; Xu, T.; Liu, X.; Yoon, J.; Xu, Z. A H-Bond Strategy to Develop Acid-Resistant Photoswitchable Rhodamine Spirolactams for Super-Resolution Single-Molecule Localization Microscopy. Chem. Sci. 2019, 10 (18), 4914–4922.

(29) Qiao, Q.; Liu, W.; Chen, J.; Wu, X.; Deng, F.; Fang, X.; Xu, N.; Zhou, W.; Wu, S.; Yin, W.; Liu, X.; Xu, Z. An Acid-Regulated Self-Blinking Fluorescent Probe for Resolving Whole-Cell Lysosomes with Long-Term Nanoscopy. Angew. Chemie Int. Ed. 2022, 61 (21), e202202961.

(30) Zheng, Y.; Ye, Z.; Zhang, X.; Xiao, Y. Recruiting Rate Determines the Blinking Propensity of Rhodamine Fluorophores for Super-Resolution Imaging. J. Am. Chem. Soc. 2023, 145 (9), 5125–5133.

(31) Zheng, Q.; Ayala, A. X.; Chung, I.; Weigel, A. V.; Ranjan, A.; Falco, N.; Grimm, J. B.; Tkachuk, A. N.; Wu, C.; Lippincott-Schwartz, J.; Singer, R. H.; Lavis, L. D. Rational Design of Fluorogenic and Spontaneously Blinking Labels for Super-Resolution Imaging. ACS Cent. Sci. 2019, 5 (9), 1602–1613.

(32) Uno, S. N.; Kamiya, M.; Morozumi, A.; Urano, Y. A Green-Light-Emitting, Spontaneously Blinking Fluorophore Based on Intramolecular Spirocyclization for Dual-Colour Super-Resolution Imaging. Chem. Commun. 2017, 54 (1), 102–105.

(33) Morozumi, A.; Kamiya, M.; Uno, S.; Umezawa, K.; Kojima, R.; Yoshihara, T.; Tobita, S.; Urano, Y. Spontaneously Blinking Fluorophores Based on Nucleophilic Addition/Dissociation of Intracellular Glutathione for Live-Cell Super-Resolution Imaging. J. Am. Chem. Soc. 2020, 142 (21), 9625–9633.

(34) Tyson, J.; Hu, K.; Zheng, S.; Kidd, P.; Dadina, N.; Chu, L.; Toomre, D.; Bewersdorf, J.; Schepartz, A. Extremely Bright, Near-IR Emitting Spontaneously Blinking Fluorophores Enable Ratiometric Multicolor Nanoscopy in Live Cells. ACS Cent. Sci. 2021, 7 (8), 1419–1426.

(35) Rahm, M.; Hoffmann, R.; Ashcroft, N. W. Atomic and Ionic Radii of Elements 1–96. Chem. - A Eur. J. 2016, 22 (41), 14625–14632.

(36) Huang, B.; Wang, W.; Bates, M.; Zhuang, X. Three-Dimensional Super-Resolution Reconstruction Microscopy. Science (80-.). 2008, 319 (5864), 810–813.

(37) Xu, F.; Ma, D.; Macpherson, K. P.; Liu, S.; Bu, Y.; Wang, Y.; Tang, Y.; Bi, C.; Kwok, T.; Chubykin, A. A.; Yin, P.; Calve, S.; Landreth, G. E.; Huang, F. Three-Dimensional Nanoscopy of Whole Cells and Tissues with in Situ Point Spread Function Retrieval. Nat. Methods 2020, 17, 531–540.

(38) Lee, M. K.; Rai, P.; Williams, J.; Twieg, R. J.; Moerner, W. E. Small-Molecule Labeling of Live Cell Surfaces for Three-Dimensional Super-Resolution Microscopy. J. Am. Chem. Soc. 2014, 136 (40), 14003–14006.

(39) Deschout, H.; Zanacchi, F. C.; Mlodzianoski, M.; Diaspro, A.; Bewersdorf, J.; Hess, S. T.; Braeckmans, K. Precisely and Accurately Localizing Single Emitters in Fluorescence Microscopy. Nat. Methods 2014, 11 (3), 253–266.

(40) Thompson, R. E.; Larson, D. R.; Webb, W. W. Precise Nanometer Localization Analysis for Individual Fluorescent Probes. Biophys. J. 2002, 82 (5), 2775–2783.

(41) Jones, S. A.; Shim, S. H.; He, J.; Zhuang, X. Fast, Three-Dimensional Super-Resolution Imaging of Live Cells. Nat. Methods 2011, 8 (6), 499–505.

(42) Bates, M.; Dempsey, G. T.; Chen, K. H.; Zhuang, X. Multicolor Super-Resolution Fluorescence Imaging via Multi-Parameter Fluorophore Detection. ChemPhysChem 2012, 13 (1), 99–107.

(43) Wang, B.; Xiong, M.; Susanto, J.; Li, X.; Leung, W. Y.; Xu, K. Transforming Rhodamine Dyes for (d)STORM Super-Resolution Microscopy via 1,3-Disubstituted Imidazolium Substitution. Angew. Chemie - Int. Ed. 2022, 61 (9), e202113612.

(44) Németh, E.; Knorr, G.; Németh, K.; Kele, P. A Bioorthogonally Applicable, Fluorogenic, Large Stokes-Shift Probe for Intracellular Super-Resolution Imaging of Proteins. Biomolecules 2020, 10 (3), 397.

(45) Man, H.; Bian, H.; Zhang, X.; Wang, C.; Huang, Z.; Yan, Y.; Ye, Z.; Xiao, Y. Hybrid Labeling System for DSTORM Imaging of Endoplasmic Reticulum for Uncovering Ultrastructural Transformations under Stress Conditions. Biosens. Bioelectron. 2021, 189 (February), 113378.

(46) Bian, H.; Ma, D.; Pan, F.; Zhang, X.; Xin, K.; Zhang, X.; Yang, Y.; Peng, X.; Xiao, Y. Cardiolipin-Targeted NIR-II Fluorophore Causes “Avalanche Effects” for Re-Engaging Cancer Apoptosis and Inhibiting Metastasis. J. Am. Chem. Soc. 2022, 144 (49), 22562–22573.

(47) Oberst, A.; Bender, C.; Green, D. R. Living with Death: The Evolution of the Mitochondrial Pathway of Apoptosis in Animals. Cell Death Differ. 2008, 15 (7), 1139–1146.

(48) Colin, J.; Gaumer, S.; Guenal, I.; Mignotte, B. Mitochondria, Bcl-2 Family Proteins and Apoptosomes: Of Worms, Flies and Men. Front. Biosci. 2009, 14 (11), 4127–4137.

(49) Hoppins, S.; Lackner, L.; Nunnari, J. The Machines That Divide and Fuse Mitochondria. Annu. Rev. Biochem. 2007, 76, 751–780.

