## Supporting Information for "Tailoring the self-blinking of sulfonamide rhodamine for long-term protein-localizing super-resolution imaging"

### 1 Experimental method

#### 1.1 General

All regents, e.g. methanol (MeOH), N,N-dimethylformamide, dichloromethane (DCM), tetrahydrofuran, triethylamine, etc., were purchased from commercial suppliers and used as received. Column chromatography was performed with silica gel (200-300 mesh and 300-400 mesh).

$^{1}H$ NMR and $^{13}C$ NMR were measured on Bruker Avance II 400, Bruker AVANCE III 500 and Varian MERCURY 400 spectrometers. Mass spectra and high-resolution mass spectra were recorded on HP 1100 LC-MSD, gas chromatography/TOF Mass, thermo Scientific LTQ Orbitrap XL, Waters Synapt G2-Si HDMS and UPLC/Q-TOF Mass spectrometers.

#### 1.2 General synthetic route

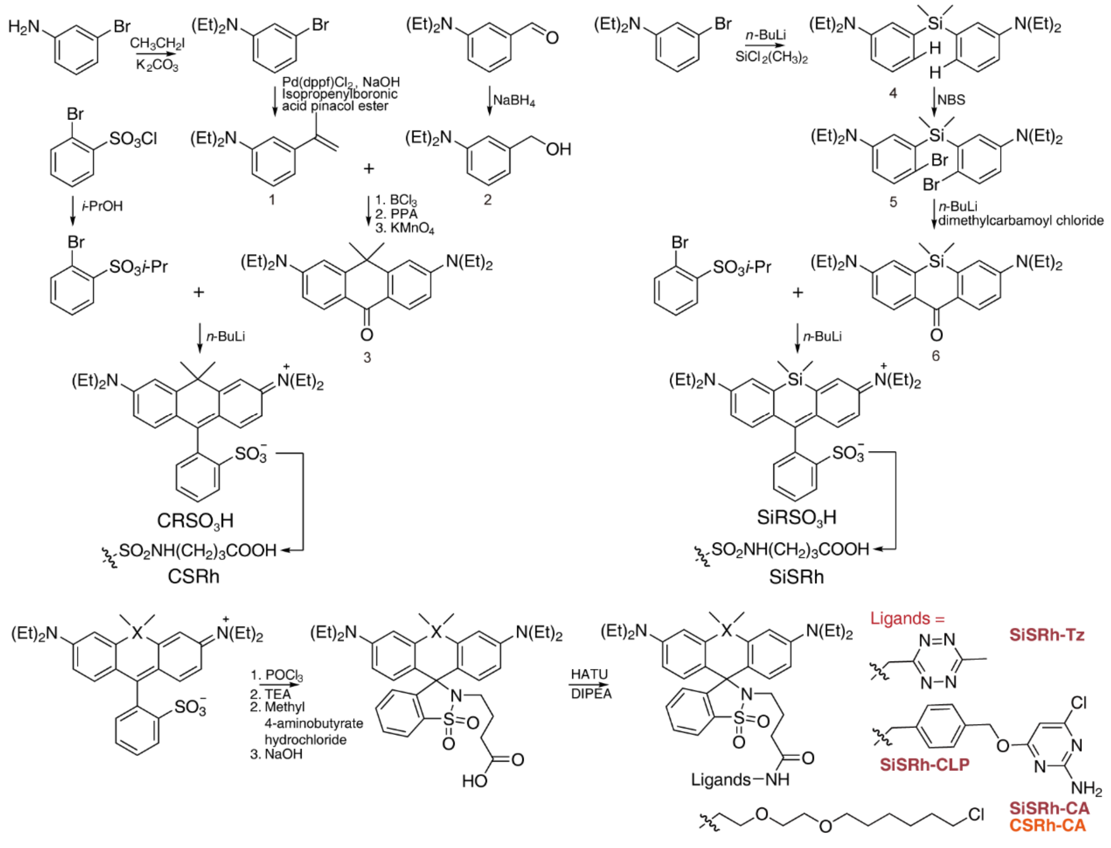

Figure S1. Synthetic routes of rhodamines.

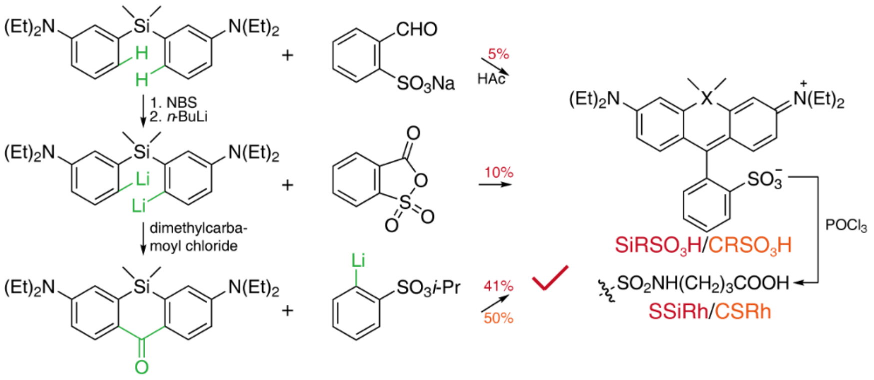

Figure S2. Comparison of the routes for the synthesis of silicon/carbon sulforhodamine.

##### 1.2.1 **General synthesis of Carbon sulforhodamine**

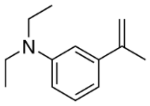

N,N-diethyl-3-(prop-1-en-2-yl)aniline (**1**): 3-bromo-N,N-diethylaniline (4 g, 17.53 mmol), potassium isopropenyltrifluoroborate (4.42 g, 26.30 mmol) and Pd(dppf)$Cl_2$ (384.89 mg, 0.53 mmol) was dissolved in 1,4-dioxane (50 mL) and filled with argon. Then, NaOH (1.4 g, 35.07 mmol) in 8 mL $H_2O$ was injected. The bottle was evacuated/backfilled with nitrogen (3×). The reaction was stirred at 80 °C for 3 h. It was then cooled to room temperature, diluted with water and extracted with EtOAc (3×). Column chromatography (0-2% EtOAc/hexanes) afforded **1** as a white solid (3.2 g, 96%). $^1H$ NMR (400 MHz, Chloroform-*d*) δ 7.16 (d, J = 7.9 Hz, 1H), 6.76 (d, J = 8.3 Hz, 2H), 6.67 – 6.58 (m, 1H), 5.31 (s, 1H), 5.04 (s, 1H), 3.37 (q, J = 7.1 Hz, 4H), 2.15 (d, J = 1.2 Hz, 3H), 1.17 (t, J = 7.0 Hz, 6H).

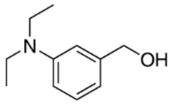

(4-(diethylamino)phenyl)methanol (**2**): 4-(diethylamino)benzaldehyde (2 g, 11.28 mmol) was dissolved in 50 mL of MeOH and $NaBH_4$ (426.88 mg, 11.28 mmol) was added slowly at 0 °C over the course of 30 minutes. Then, the mixture was stirred for 1 hour at room tmeperature. Reaction was quenched of water and solvent was evaporated on rotary evaporator. The residue was dissolved with water and extracted with EtOAc, washed with brined, dried over $Na_2SO_4$ and filtered. The solvent was evaporated and product was purified by flash column chromatography (20% EtOAc/hexanes) to obtained **2** as a white solid (1.6 g, 79%). The product was stored under inert gas at -20 °C for direct used in the next step.

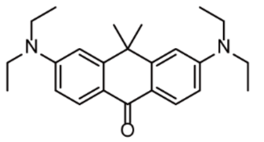

3,6-bis(diethylamino)-10,10-dimethylanthracen-9(10H)-one (**3**): $BCl_3$ (2.35 mL, 2.35 mmol, 1 M) was slowly added to dry DCM (5 mL) containing compounds 1[1] (500 mg, 1.96 mmol) and 2[1] (351 mg, 1.96 mmol) under argon at -78 °C. After stirring for a few minutes, the reaction solution slowly rose to room temperature and was stirred overnight. The reaction solution was directly evaporated to produce crude residue. The crude residue was dissolved in polyphosphate (7 g) and reacted at 110 °C for 5 h. The end point was determined by TLC. The reaction solution was poured into ice, neutralized by NaOH, extracted by DCM, washed by salt water, dried by anhydrous sodium sulfate, and the crude residue was obtained by spin drying. The crude residue was dissolved in acetone (15 mL), decreased to -15 °C, and then added in small batches of powdered $KMnO_4$ (740 mg) within half an hour. After addition, the reaction continued for 2 h, TLC determined the end point, and then the reaction liquid was poured into DCM (15 mL) at -78 °C to quench the reaction. The solution is then filtered on funnel lined with diatomaceous earth and purified by the column to obtain the yellow product **3** (620 mg, 65%). $^1H$ NMR (400 MHz, Chloroform-*d*) δ 8.23 (d, J = 9.0 Hz, 2H), 6.72 (d, J = 7.3 Hz, 4H), 3.47 (q, J = 7.1 Hz, 8H), 1.70 (s, 6H), 1.25 (t, J = 7.0 Hz, 12H).

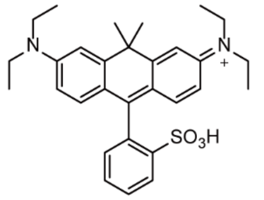

Carbon sulforhodamine (**4**): propan-2-yl 2-bromobenzene-1-sulfonate (1.23 g, 8 eq) was dissolved in dry tetrahydrofuran (12 mL) under argon. The solution was cooled to -20 °C, and *n*-BuLi (2.5 M in hexane, 1.65 mL, 7.5 eq) was added slowly. After 10 min, dissolved 3 (200 mg, 1 eq) in 3 ml dry tetrahydrofuran and added it dropwise to the reaction mixture, stirring at rt for 1 h. The solvent was removed by rotary evaporation, and the crude product was further purified by silica column (DCM:MeOH = 20:1) to give Carbon Sulforhodamine (**4**) as a blue solid (145 mg, 52%). $^1H$ NMR (400 MHz, DMSO-$d_6$) δ 7.94 (dd, J = 7.9, 1.4 Hz, 1H), 7.56 (td, J = 7.6, 1.4 Hz, 1H), 7.48 (td, J = 7.4, 1.4 Hz, 1H), 7.13 (d, J = 2.5 Hz, 2H), 7.08 (dd, J = 7.5, 1.3 Hz, 1H), 6.92 (d, J = 9.4 Hz, 2H), 6.79 (dd, J = 9.5, 2.4 Hz, 2H), 3.66 (q, J = 7.2 Hz, 8H), 1.80 (s, 3H), 1.67 (s, 3H), 1.20 (t, J = 7.0 Hz, 12H). $^{13}C$ NMR (101 MHz, DMSO-$d_6$) δ 166.48, 156.37, 153.92, 147.07, 138.25, 132.35, 129.42, 128.66, 128.21, 127.81, 120.70, 112.05,

109.99, 45.12, 41.30, 34.88, 31.20, 12.80. HRMS *m/z*: calcd for $C_{30}H_{37}N_2O_3S^+$ $[M]^+$ 505.2525; found: 505.2516.

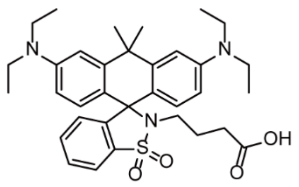

CSRh-COOH: Carbon Sulforhodamine (**4**, 50 mg, 1 eq) was dissolved in dry 1,2-dichloroethane (5 mL) under argon. Phosphorus oxychloride (16.68 mg, 1.1 eq) was added at room temperature for 5 min. Then the solution was refluxed for 2 h. After cooling to room temperature, the mixture of crude acid chloride was added to a DCM/MeCN (5mL/5 mL) mixture solution of 4-amino-butyricacimethylester (18 mg, 1.2 eq) and triethylamine at ice bath. After stirring 2h, the crude product was further purified by silica column (petroleum ether: ethylene acetate = 2:1) to give CSRh-COOMe as a colorless solid. CSRh-COOMe (1 eq) and NaOH (19 mg, 5 eq) was dissolved in methane/water (4 mL/1 mL) and refluxed 3h. After completion of the reaction, the solution was removed by rotary evaporation, and the crude product was further purified by silica column (DCM: MeOH = 30:1) to give CSRh-COOH as a light blue solid (85%). $^1$H NMR (500 MHz, DMSO-$d_6$) δ 7.92 (d, J = 7.4 Hz, 1H), 7.50 (t, J = 7.4 Hz, 2H), 6.81 – 6.68 (m, 5H), 6.55 (d, J = 8.9 Hz, 2H), 3.40 – 3.33 (m, 8H), 2.78 (t, J = 7.3 Hz, 2H), 1.98 (t, J = 7.4 Hz, 2H), 1.78 (s, 3H), 1.71 (s, 3H), 1.39 (t, J = 7.3 Hz, 2H), 1.09 (t, J = 7.0 Hz, 12H). $^{13}$C NMR (101 MHz, DMSO-d$_6$) δ 173.72, 146.89, 146.81, 145.37, 133.69, 131.96, 129.17, 128.81, 125.93, 120.16, 118.70, 111.61, 108.10, 69.39, 43.55, 36.97, 35.08, 34.44, 30.87, 23.66, 12.52. HRMS *m/z*: calcd for $C_{34}H_{43}N_3O_4S$ $[M+H]^+$ 590.3008; found: 590.3034.

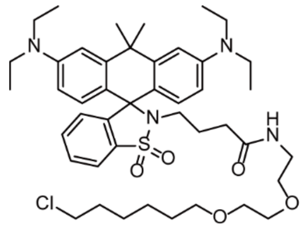

CSRh-CA: CSRh-COOH (20 mg, 1 eq), Halo-$NH_2$ (9 mg, 1.2 eq), and HATU (15 mg, 1.2 eq) were dissolved in dry N,N-dimethylformamide (2 mL) in the presence of N,N-disopropylethylamine (21.9 mg, 5 eq). The mixture was stirred at room temperature for 2 h. The reaction mixture was further washed with brine, dried over $Na_2SO_4$, filtered and evaporated. The crude product was further purified by silica column (DCM: MeOH = 40:1) to give CSRh-CA as a light blue solid (60%). $^1$H NMR

(400 MHz, DMSO-$d_6$) δ 7.91 (d, J = 7.4 Hz, 1H), 7.54 (t, J = 5.9 Hz, 1H), 7.48 (q, J = 6.8 Hz, 2H), 6.75 (t, J = 7.2 Hz, 2H), 6.71 (d, J = 3.0 Hz, 2H), 6.55 (dd, J = 9.0, 2.4 Hz, 2H), 3.60 (t, J = 6.6 Hz, 2H), 3.42 (s, 4H), 3.35 (d, J = 4.6 Hz, 8H), 3.26 (t, J = 6.1 Hz, 2H), 3.03 (q, J = 5.9 Hz, 2H), 2.74 (t, J = 7.8 Hz, 2H), 1.99 (p, J = 7.0, 6.4 Hz, 2H), 1.84 (t, J = 7.4 Hz, 2H), 1.79 (s, 3H), 1.70 (d, J = 6.1 Hz, 5H), 1.47 – 1.29 (m, 8H), 1.09 (t, J = 6.9 Hz, 12H). $^{13}$C NMR (101 MHz, DMSO-$d_6$) δ 171.20, 146.87, 145.39, 133.67, 131.96, 129.64, 129.19, 125.92, 120.16, 118.75, 111.56, 108.06, 70.16, 69.53, 69.38, 68.99, 45.34, 43.54, 38.33, 36.99, 35.00, 34.58, 32.71, 26.09, 24.91, 24.74, 12.56. HRMS *m/z*: calcd for $C_{44}H_{63}ClN_4O_5S$ $[M+H]^+$ 795.4241; found: 795.4278.

1.2.2 **General synthesis of Silicon Sulforhodamines**

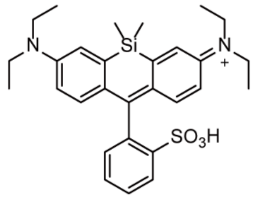

Silicon Sulforhodamine (**8**):

Route A: 5[2] (200 mg, 1 eq) and 2-formylbenzenesulfonic acid sodium salt (4 eq) was dissolved in 3 ml AcOH and stirred at 170 °C for 8 h. After cooling to room temperature, the solvent was removed by rotary evaporation, and the crude product was further purified by silica column (DCM:MeOH = 20:1) to give Silicon Sulforhodamine (**8**) as a blue solid (5%).$^1$H NMR (400 MHz, Methanol-$d_4$) δ 8.14 (d, J = 7.7 Hz, 1H), 7.61 (dt, J = 22.3, 7.4 Hz, 2H), 7.23 (d, J = 2.6 Hz, 2H), 7.16 (d, J = 7.4 Hz, 1H), 7.07 (d, J = 9.6 Hz, 2H), 6.69 (dd, J = 9.6, 2.6 Hz, 2H), 3.68 (q, J = 7.1 Hz, 8H), 1.27 (t, J = 7.1 Hz, 12H), 0.58 (s, 6H). $^{13}$C NMR (126 MHz, Methanol-$d_4$) δ 171.62, 153.77, 149.40, 145.45, 143.97, 137.79, 131.50, 130.66, 129.91, 129.53, 129.22, 121.16, 114.39, 46.61, 13.11,. HRMS m/z: calcd for $C_{29}H_{37}N_2O_3SSi^+$ $[M]^+$ 521.2289; found: 521.2288.

Route B: 6[2] (1g, 1 eq) was dissolved in 50 ml dry Tetrahydrofuran. The solution was cooled to -78 °C, and *t*-BuLi (1.3 M in hexane, 6.61 mL, 4.4 eq) was added slowly. After 40 min, dissolved 2-Sulfobenzoic anhydride (899 mg, 2.5 eq) in 20 ml dry Tetrahydrofuran and added it dropwise to the reaction mixture, stirring at rt for 1 h. The solvent was removed by rotary evaporation, and the crude product was further purified by silica column (DCM: MeOH = 20:1) to give Silicon Sulforhodamine (**8**) as a blue solid (10%).

Route C: Propan-2-yl 2-bromobenzene-1-sulfonate (5.86 g, 8 eq) was dissolved in 50 ml dry tetrahydrofuran. The solution was cooled to -20 °C, and *n*-BuLi (2.5 M in hexane, 7.8 mL, 7.5 eq) was added slowly. After 10 min, dissolved 7[2] (1 g, 1 eq) in 5 ml dry tetrahydrofuran and added it dropwise to the reaction mixture, stirring at rt for 1 h.

The solvent was removed by rotary evaporation, and the crude product was further purified by silica column (DCM:MeOH = 20:1) to give Silicon Sulforhodamine (**8**) as a blue solid (45%).

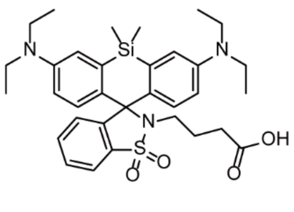

SiSRh-COOH: Silicon Sulforhodamine (**8**, 100 mg, 1 eq) was dissolved in dry 1,2-dichloroethane (5 mL) under argon. Phosphorus oxychloride (32.39 mg, 1.1 eq) was added at room temperature for 5 min. Then the solution was refluxed for 2 h. After cooling to room temperature, the mixture of crude acid chloride was added to a DCM/MeCN (5 mL/5 mL) mixture solution of 4-amino-butyricacimethylester (1.1 eq) and triethylamine at ice bath. After stirring 2h, the crude product was further purified by silica column (petroleum ether: ethylene acetate = 2:1) to give SiSRh-COOMe as a colorless solid. SiSRh-COOMe (1 eq) and NaOH (5 eq) was dissolved in methane/$H_2O$ (8 mL/2mL) and refluxed 3h. After completion of the reaction, the solution was removed by rotary evaporation, and the crude product was further purified by silica column (DCM: MeOH = 30:1) to give SiSRh-COOH as a light blue solid (85%). $^1H$ NMR (500 MHz, Chloroform-*d*) δ 7.84 (d, J = 7.6 Hz, 1H), 7.36 (dt, J = 21.9, 7.3 Hz, 2H), 7.13 (d, J = 9.1 Hz, 2H), 6.79 (d, J = 7.6 Hz, 1H), 6.72 (s, 2H), 6.64 (d, J = 9.0 Hz, 2H), 3.34 (q, J = 6.9 Hz, 8H), 2.98 (t, J = 6.8 Hz, 2H), 2.16 (t, J = 7.3 Hz, 2H), 1.69 – 1.61 (m, 2H), 1.15 (t, J = 7.0 Hz, 12H), 0.53 (s, 3H), 0.48 (s, 3H). HRMS m/z: calcd for $C_{33}H_{44}N_3O_4SSi$ $[M+H]^+$ 606.2816; found: 606.2819.

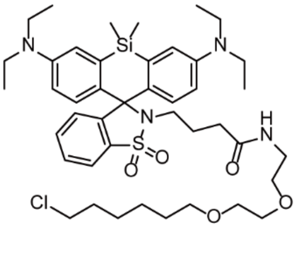

SiSRh-CA: SiSRh-COOH (20 mg, 1 eq), Halo-$NH_2$ (8.86 mg, 1.2 eq), and HATU (15 mg, 1.2 eq) were dissolved in dry N,N-dimethylformamide (2 mL) in the presence of N,N-disopropylethylamine (21 mg, 5eq). The mixture was stirred at room temperature for 2 h. The reaction mixture was further washed with brine, dried over $Na_2SO_4$, filtered and evaporated. The crude product was further purified by silica column (DCM: MeOH = 30:1) to give SiSRh as a light blue solid (76%). $^1H$ NMR (500 MHz, Chloroform-*d*) δ 7.83 (d, J = 7.6 Hz, 1H), 7.37 (dt, J = 22.7, 7.3 Hz, 2H), 7.11 (d,

J = 9.1 Hz, 2H), 6.78 (d, J = 7.6 Hz, 1H), 6.72 (d, J = 2.8 Hz, 2H), 6.63 (dd, J = 9.1, 2.8 Hz, 2H), 3.62 – 3.56 (m, 4H), 3.50 (ddd, J = 18.4, 13.3, 6.7 Hz, 6H), 3.41 – 3.28 (m, 10H), 2.95 (t, J = 6.3 Hz, 2H), 2.00 (t, J = 7.1 Hz, 2H), 1.76 (dd, J = 14.6, 6.9 Hz, 2H), 1.61 (d, J = 6.4 Hz, 4H), 1.44 (dd, J = 15.0, 7.8 Hz, 2H), 1.38 (dd, J = 15.0, 8.1 Hz, 2H), 1.16 (t, J = 7.0 Hz, 12H), 0.53 (s, 3H), 0.48 (s, 3H). $^{13}$C NMR (126 MHz, Chloroform-*d*) δ 146.12, 133.49, 131.90, 131.54, 128.31, 125.89, 120.76, 114.63, 113.96, 71.36, 70.41, 70.16, 69.91, 45.15, 44.23, 40.75, 39.26, 34.11, 32.67, 29.60, 26.82, 25.54, 24.64, 12.73. HRMS m/z: calcd for $C_{43}H_{63}ClN_4O_5SSi$ $[M+H]^+$ 811.4011; found: 811.4044.

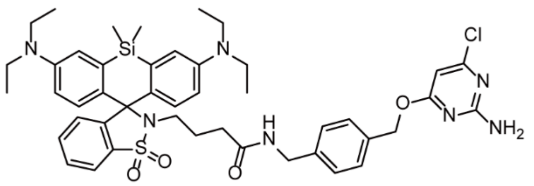

SiSRh-CLP: To a solution of SiSRh-COOH (20 mg, 1 eq) in 5 ml DMSO were successively added HATU (18.7 mg, 1.5 eq) and N,N-disopropylethylamine (21 mg, 5eq). After 1 min, CLP-$NH_2$ (13 mg, 1.5 eq) were added. The mixture was stirred at room temperature for 2 h. The reaction mixture was further washed with brine, dried over $Na_2SO_4$, filtered and evaporated. The crude product was further purified by silica column (DCM: MeOH = 30:1) to give SiSRh-CLP as a light blue solid (53%). $^1$H NMR (400 MHz, DMSO-$d_6$) δ 7.89 (dd, J = 6.2, 2.8 Hz, 1H), 7.46 (dd, J = 5.8, 3.1 Hz, 2H), 7.32 (d, J = 8.1 Hz, 2H), 7.14 (d, J = 7.9 Hz, 2H), 7.04 (d, J = 9.1 Hz, 2H), 6.78 (d, J = 2.9 Hz, 2H), 6.73 – 6.69 (m, 1H), 6.67 (dd, J = 9.2, 2.9 Hz, 2H), 6.12 (s, 1H), 5.27 (s, 2H), 4.12 (d, J = 5.8 Hz, 2H), 3.35 (d, J = 7.1 Hz, 8H), 2.88 – 2.80 (m, 2H), 1.92 (t, J = 7.4 Hz, 2H), 1.57 (p, J = 7.6 Hz, 2H), 1.07 (t, J = 6.9 Hz, 12H), 0.55 (s, 3H), 0.47 (s, 3H). $^{13}$C NMR (126 MHz, DMSO-$d_6$) δ 171.10, 170.25, 162.74, 159.96, 146.95, 145.51, 139.43, 134.50, 134.24, 133.62, 131.17, 131.00, 130.13, 128.40, 128.23, 127.10, 120.47, 114.06, 113.86, 94.35, 73.79, 43.31, 41.69, 40.70, 32.91, 24.63, 12.41. HRMS m/z: calcd for $C_{45}H_{54}ClN_7O_4SSi$ $[M+H]^+$ 852.3449; found: 852.3457

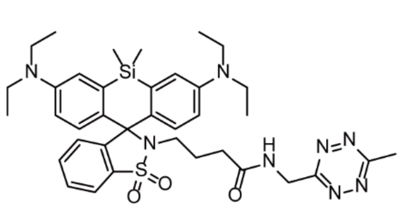

SiSRh-Tz: SiSRh-COOH (20 mg, 1 eq), Tz-$NH_2$ (4.96 mg, 1.2 eq), and HATU (15 mg, 1.2 eq) were dissolved in 5 ml DCM in the presence of N,N-disopropylethylamine (21 mg, 5eq). The mixture was stirred at room temperature for 2 h. The reaction mixture was further washed with brine, dried over $Na_2SO_4$, filtered and evaporated. The crude product was further purified by silica column (DCM: MeOH = 30:1) to give SiSRh-Tz as a light blue solid (42%). $^1$H NMR (500 MHz, Chloroform-*d*) δ 7.84 (d, J = 7.7 Hz, 1H), 7.38 (dt, J = 22.3, 7.2 Hz, 2H), 7.10 (d, J = 9.0 Hz, 2H), 6.82 – 6.61 (m, 5H), 4.89 (d, J = 5.7 Hz, 2H), 3.32 (q, J = 6.9 Hz, 8H), 3.03 (d, J = 7.0 Hz, 5H), 2.15 (t, J = 6.8 Hz, 2H), 1.69 – 1.61 (m, 2H), 1.14 (t, J = 7.0 Hz, 12H), 0.51 (d, J = 23.4 Hz, 6H). HRMS m/z: calcd for $C_{37}H_{49}N_8O_3SSi$ $[M+H]^+$ 713.3412; found: 713.3409.

#### 2 **Spectroscopic study**

##### 2.1 **Method**

Absorption spectra were recorded on Agilent 8453 UV-visible spectrophotometer and fluorescence spectra were recorded on Agilent Cary Eclipse Fluorescence spectrophotometer.

###### 2.1.1 **p$K_{cycling}$ measurement.**

Unbounded fluorophore: Fluorophores were prepared as 2 μM solution for measurements. The buffer was PBS containing 30% ethanol. The proton responding curve was obtained through pH titration of the fluorophore solution.

Fluorophore Halo-protein conjugates: Fluorophores were labeled to Halo-protein in PBS (pH=7.4) at 37°C for 2 h. The remaining unconjugated fluorophores were removed by protein desalting spin columns (filled with Sephadex G-25 resins). Next, fluorescence spectra were measured in PBS at various pH values.

PBS buffer was 100 mM phosphate buffered saline.

*p*Ka was calculated through Henderson–Hasselbalch equation,

$$I = \frac{1}{b(10^{pH-pKa} + 1)} + c$$

*I* was the integration of fluorescence peak or absorption peak. *b* and *c* were constants.

###### 2.1.2 **Equilibrium rates measurement**

Unbounded fluorophore: Fluorophores were prepared as 2 μM solution in MilliQ water containing 30% EtOH for measurements. A mixed buffer was rapidly changed from an acidic environment around pH 4 to an alkaline environment at pH 10 to detect the change of the fluorescence intensity with time. Another mixed buffer was rapidly changed from pH 10 to 4 in an acidic environment to detect the fluorescence intensity as a function of time.

Fluorophore Halo-protein conjugates: First, fluorophores were labeled to Halo-protein in PBS (pH=7.4) by incubated at 37°C for 2h. The remaining unbounded fluorophores were removed by protein desalting spin columns (filled with Sephadex

G-25 resins). Next, the fluorophore-protein were prepared in PBS buffer for measurements. The method is the same as the unbounded fluorophore.

2.2 **Analysis**

The kinetics of intramolecular spirocyclization of CSRh and SiSRh were tested in vitro through spectroscopy, and the fluorescence response of the ring opening and closing were recorded by changing the pH of the system (Figure S1). After adding acids, the free fluorophores rapidly changed their equilibrium within 20 seconds, showing fast acid equilibrium rates ($k_{E\text{-}a}$ = 0.24 $s^{-1}$, SiSRh; $k_{E\text{-}a}$ = 0.15 $s^{-1}$, CSRh). However, when the fluorophores bind to Halo protein and acid is added, it is found that the time to reach equilibrium slows down, exhibiting slower acid equilibrium rates ($k_{E\text{-}a}$ = 0.03 $s^{-1}$, SiSRh + protein; $k_{E\text{-}a}$ = 0.05 $s^{-1}$, CSRh + protein). After adding bases, both the free fluorophores and the fluorophore Halo-protein conjugates can quickly switch fluorescence states and reach equilibrium ($k_{E\text{-}b}$ = 0.34 $s^{-1}$, SiSRh; $k_{E\text{-}b}$ = 0.31 $s^{-1}$, CSRh; $k_{E\text{-}b}$ = 0.34 $s^{-1}$, SiSRh + protein; $k_{E\text{-}b}$ = 0.30 $s^{-1}$, CSRh + protein). Both the free fluorophore, as well as the fluorophore protein conjugates of SiSRh and CSRh, exhibit slower acid equilibrium rates compared to HMSiR and SRhB. This indicates that sulfonamide groups and more electrophilic xanthene cores can slow down the ring opening rate of fluorophores, and the ring opening rate provides a new method for studying the trend of imaging time of fluorophores.

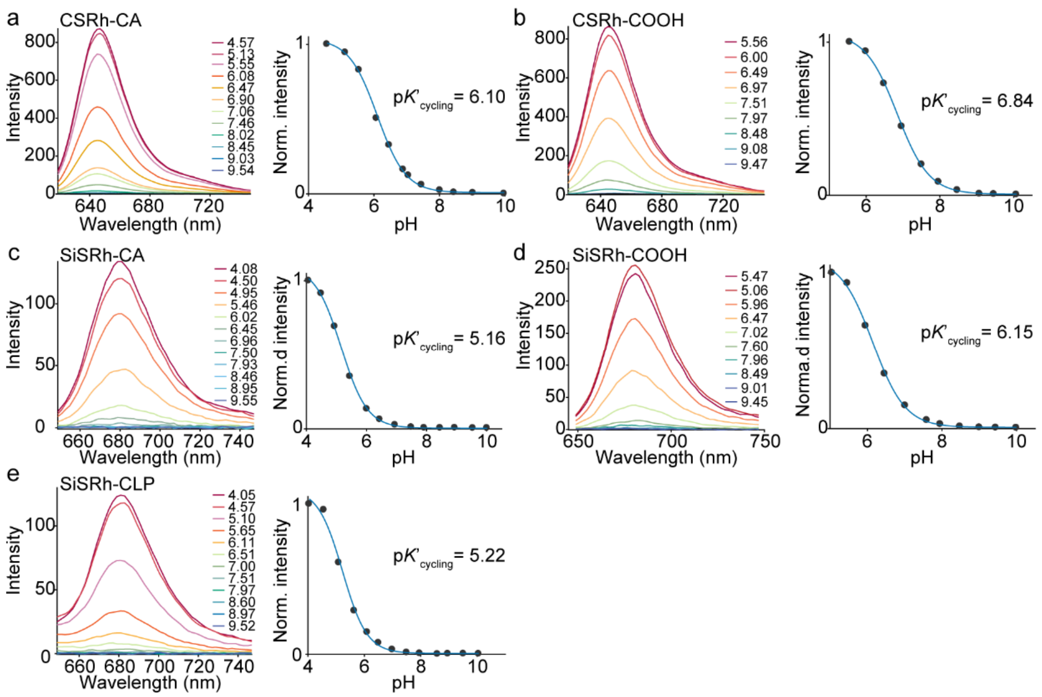

Figure S3. pH titration of silicon sulfonamide rhodamine and carbon sulfonamide rhodamines. The titration was performed with free fluorophore molecules in PBS/EtOH (v/v = 7: 3).

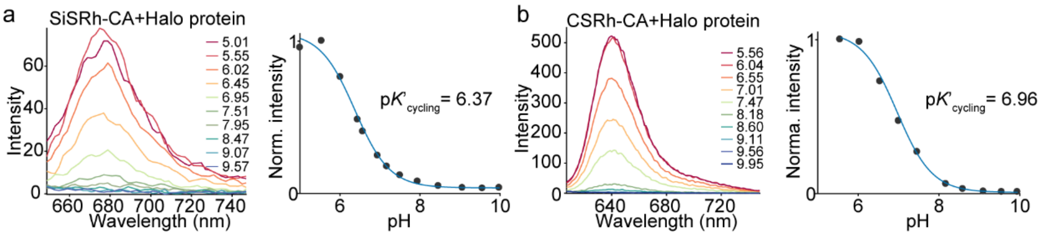

Figure S4. pH titration of sulfonamide rhodamines bounded to Halo proteins. The titration was performed with fluorophore protein conjugates in PBS. The spectra outside pH range 5.0-10.0 were not meaningful as proteins degenerated under our conditions.

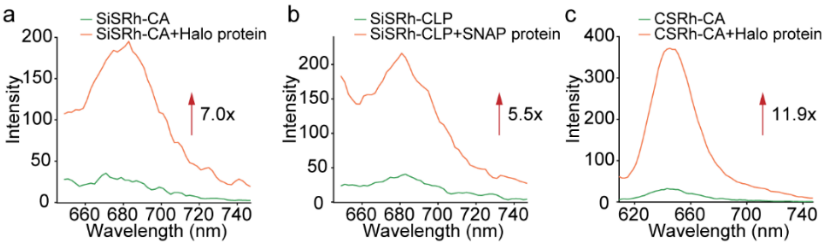

Figure S5. Fluorescent spectra of sulfonamide rhodamines in the absence (green line) and presence Halo-protein (orange line) after 2 h incubation.

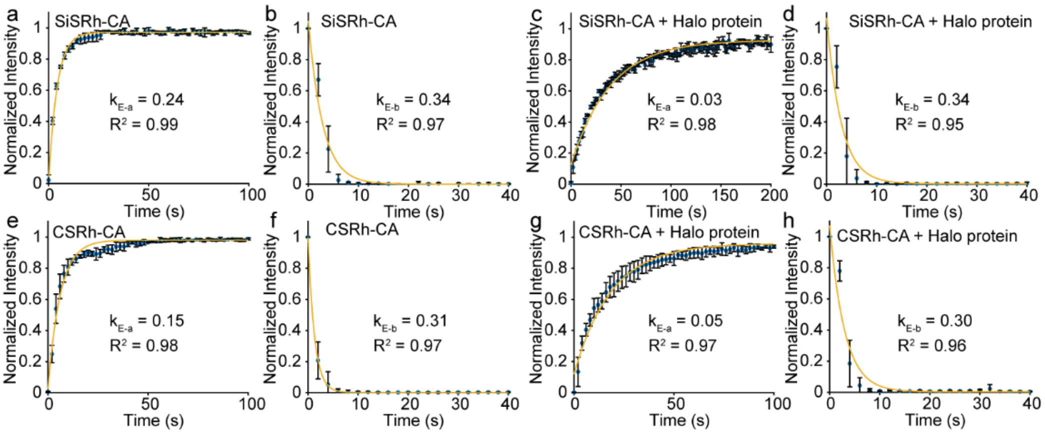

Figure S6. Ensemble kinetic study of spirocyclization equilibriums for sulfonamide rhodamines before and after labeling with Halo protein under acidic or alkaline disturbances. Peak emission intensities (680 nm for SiSRh and 640 nm for CSRh) are plotted as a function of time, showing the rates reflecting equilibrium shifts to ring-opening ($k_{E\text{-}a}$) and ringclosing states ($k_{E\text{-}b}$).

### 3 **Single-Molecule study**

#### 3.1 **Method**

Coverslips (Fisherbrand, 12-545-102) utilized in this study were cleaned through sonication in ethanol, 1 M KOH and water. The cleaned coverslips were stored in

MilliQ water. Before imaging. these coverslips were dried under dry nitrogen gas. Fluorophore was labeled to Halo protein in PBS (pH=7.4), then the resulting mixture was incubated at 37 °C for 2 h. The remaining unbounded fluorophores were removed by protein desalting spin columns (filled with Sephadex G-25 resins). The protein solution labeled with fluorophores were freshly diluted in PBS at a concentration low enough to minimize the overlapping between different molecules and transferred to the surface of clean coverslips. The adhesion of proteins to the surface proceed for 30 s and the unbounded proteins were washed out through three times rinses of PBS. The pixel size was 160 nm/pixel and 10000 raw frames were acquired at 10 ms exposure time under different laser power irradiation. At least five measurements were performed for each irradiation condition.

Single-molecule data were automatically processed with a home-written Matlab software as described in our previous report.[3, 4] Briefly, a wavelet filter was utilized to remove background noises on raw image stacks. Single-molecule candidates were identified from filtered image stacks and their single molecule fluorescent trajectories were further extracted from unfiltered raw data. These trajectories were analyzed with a hidden Markov model to obtain the state transition trajectories through Baum–Welch, forward-backward and Viterbi algorithms (A gaussian distribution model was deployed for the observing probability distribution). Single-molecule photophysics were measured from single-molecule trajectories and their state transition trajectories.

##### 3.2 **Results**

Table S1. Single-molecule Photophysics of SiSRh, CSRh and SRhB

| Dye | Laser power ($W/cm^2$) | Brightness (photons/10ms) | Duty cycle | Switch numbers | Dark time (s) | Bleach time (s) | $k_{rc}$ ($s^{-1}$) |
|---|---|---|---|---|---|---|---|
| SiSRh | 500 | 143±7 | 0.005±0.0003 | 10.7±0.3 | 5.4±0.2 | 52.6±1.0 | 0.18±0.02 |
| | 800 | 242±6 | 0.004±0.0006 | 9.2±1.1 | 5.2±0.2 | 43.0±4.7 | 0.17±0.01 |
| CSRh | 500 | 95±10 | 0.011±0.0022 | 24.9±2.8 | 2.1±0.1 | 51.4±5.1 | 0.46±0.10 |
| | 800 | 122±4 | 0.009±0.0018 | 20.0±4.0 | 2.4±0.4 | 47.0±2.2 | 0.45±0.11 |
| SRhB | 500 | 128±10 | 0.022±0.0023 | 17.1±3.2 | 0.4±0.1 | 8.9±1.0 | 2.80±0.62 |
| | 800 | 187±32 | 0.014±0.0015 | 11.4±2.0 | 0.4±0.1 | 5.6±0.6 | 3.07±0.64 |
| HMSiR | 500 | 182±11 | 0.012±0.0009 | 13.6±1.3 | 2.2±0.2 | 28.6±2.7 | 0.41±0.05 |
| | 800 | 284±5 | 0.007±0.0006 | 8.0±0.9 | 2.8±0.2 | 20.1±1.3 | 0.38±0.05 |

| | | | | | | | |
|---|---|---|---|---|---|---|---|
| AF647 | 500 | 348±25 | 0.002±0.0003 | 2.0±0.2 | 0.5±0.2 | 0.81±0.3 | 18.4±3.0 |
| | 800 | 592±37 | 0.001±0.0001 | 1.8±0.1 | 0.3±0.2 | 0.39±0.2 | 12.2±1.8 |

### 4 Super-resolution Imaging

#### 4.1 Total internal reflection microscopy

Super-resolution imaging was studied with a total internal reflection fluorescence microscope (TIRFM) built on an Olympus IX71 inverted microscope as described earlier.[3, 5] The laser light was focused on the back focal plane of a × 100 objective (UAPON 100XOTIRF; 1.49 numerical aperture). An EMCCD camera (iXon DU-897U) was implemented for data acquisition.

#### 4.2 Post-processing method

Super-resolution imaging analysis was performed in either a ThunderStorm[6] plugin of ImageJ[7] and our home-written Matlab software. Briefly, the raw frames were filtered with a difference-of-Gaussians filter to search for signal candidates. Then the point spread functions (PSF) of those candidates were fitted with an integrated form of symmetric 2D Gaussian function (Fitting radius: 3.0 pixel) following Maximum likelihood method[8, 9] to estimate the precise location and single-molecule intensity. The localization precision was calculated according to the Thompson formula.[10] Those PSFs with large widths (> 1.5×median(sigma)) or small widths (< 0.5×median(sigma)) were removed.

Camera readout intensity (I) was converted to photons through the below equations:

$$\text{photons} = \frac{\text{I} \times \text{ADE}}{\text{QE} \times \text{EMGain}}$$

I was the intensity value direct read from camera. ADU was the sensitivity of EMCCD (15.82 electrons per A/D count). EMGain was the gain configuration of the camera (100 in this study). QE was the photon efficiency of the camera.

#### 4.3 Cell culture & staining

Cell Culture. HeLa cells were purchased from the Cell Bank of Type Culture of Chinese Academy of Sciences. Vero and U2OS cells were kindly provided by Procell Life Science&Technology Co., Ltd. HeLa and Vero cells were cultured in full growth medium (cell culture media), that is, minimum Eagle's medium (MEM) supplemented with 10% fetal bovine serum (FBS, HyClone) and 1% penicillin−streptomycin (PS) solution (×100 HyClone). U2OS cells were cultured in full growth medium (cell culture media), that is, McCoy's 5A(PM150710) supplemented with 10% FBS

(HyClone) and 1% PS solution (×100 HyClone). The culturing condition was a humidified atmosphere at 37 °C charged with 5% $CO_2$. HeLa and U2OS cells were transient transfected with Halo-H2B, Halo-Sec61β (Addgene plasmid #123285), Halo-TOMM20 (Addgene plasmid #123284), SNAP-COX8A (Addgene plasmid#101129), and SNAP-ER[11] using Lipofectamine 3000 reagent following a standard protocol. Vero cells were transient transfected with Halo-βTubulin (Addgene plasmid #64691) using Lipofectamine 3000 reagent following a standard protocol. Cells were seeded in cover slips after 24 h of transfection.

Super-Resolution Imaging Acquisition. Halo-H2B expressing cells were incubated with 50 nM SiSRh-CA and 10 nM CSRh-CA. Halo-Sec61β-expressing cells were incubated with 400 nM SiSRh-CA, 300 nM SRhB, 300 nM HMSiR and 100 nM CSRh-CA. Halo-βTubulin-expressing cells were incubated with 2 μM SiSRh-CA, and 1 μM CSRh-CA. Halo-TOMM20-expressing cells were incubated with 100 nM SiSRh-CA, 100 nM HMSiR, 100 nM SRhB, and 50 nM CSRh-CA. SiSRh-CA, CSRh-CA, and SRhB were stained in cells for 2 h; HMSiR was stained in cells for 8 h. The free remaining dyes were washed with PBS for three times and were further cultured in a $CO_2$ incubator with fresh MEM or McCoy's 5A(PM150710) media for 30 min. The imaging media was MEM or McCoy's 5A(PM150710) without phenol red supplemented with 10% FBS.

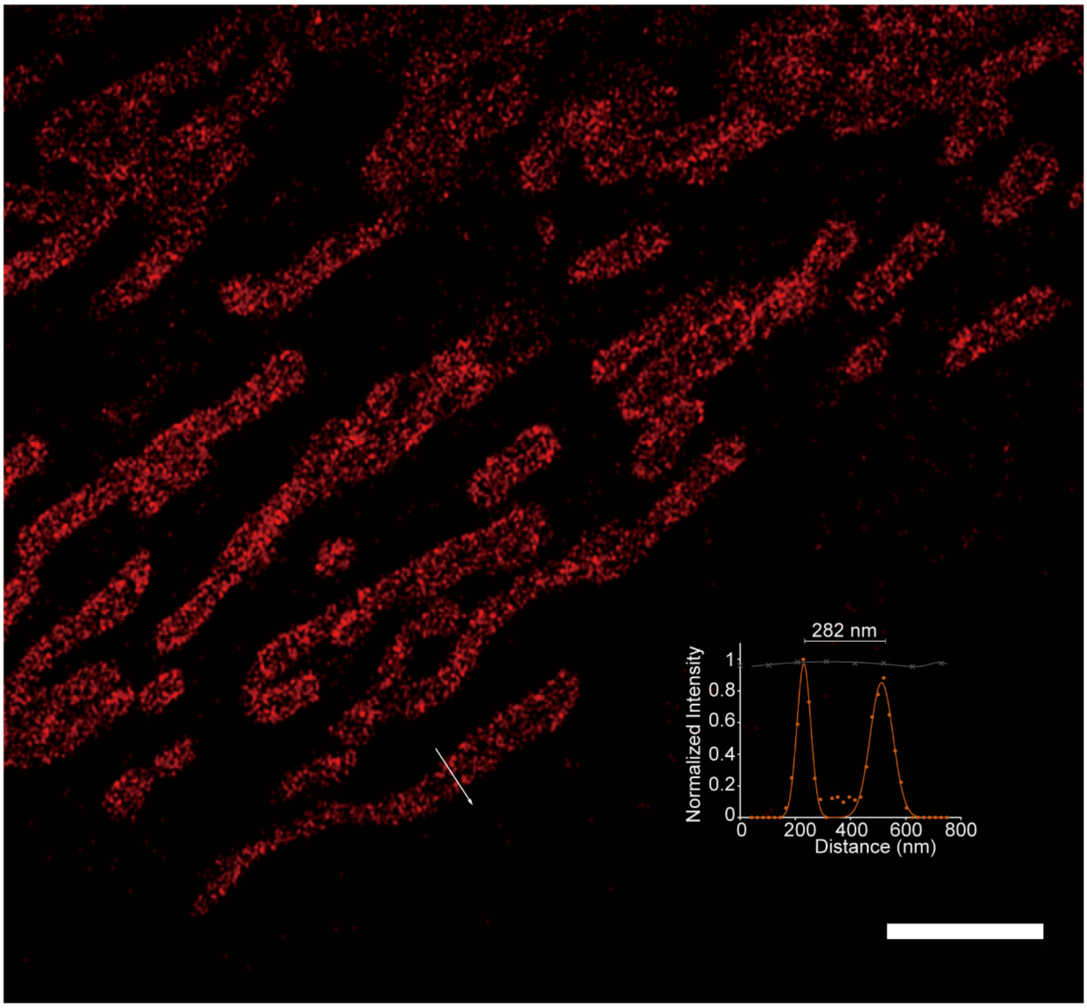

Figure S7. Super-resolution imaging of mitochondrial in living U2OS cells with SiSRh-CA. The inset represents intensity profiles of mitochondrial highlighted with white lines in the super-resolution image. Scale bars: 2 μm.

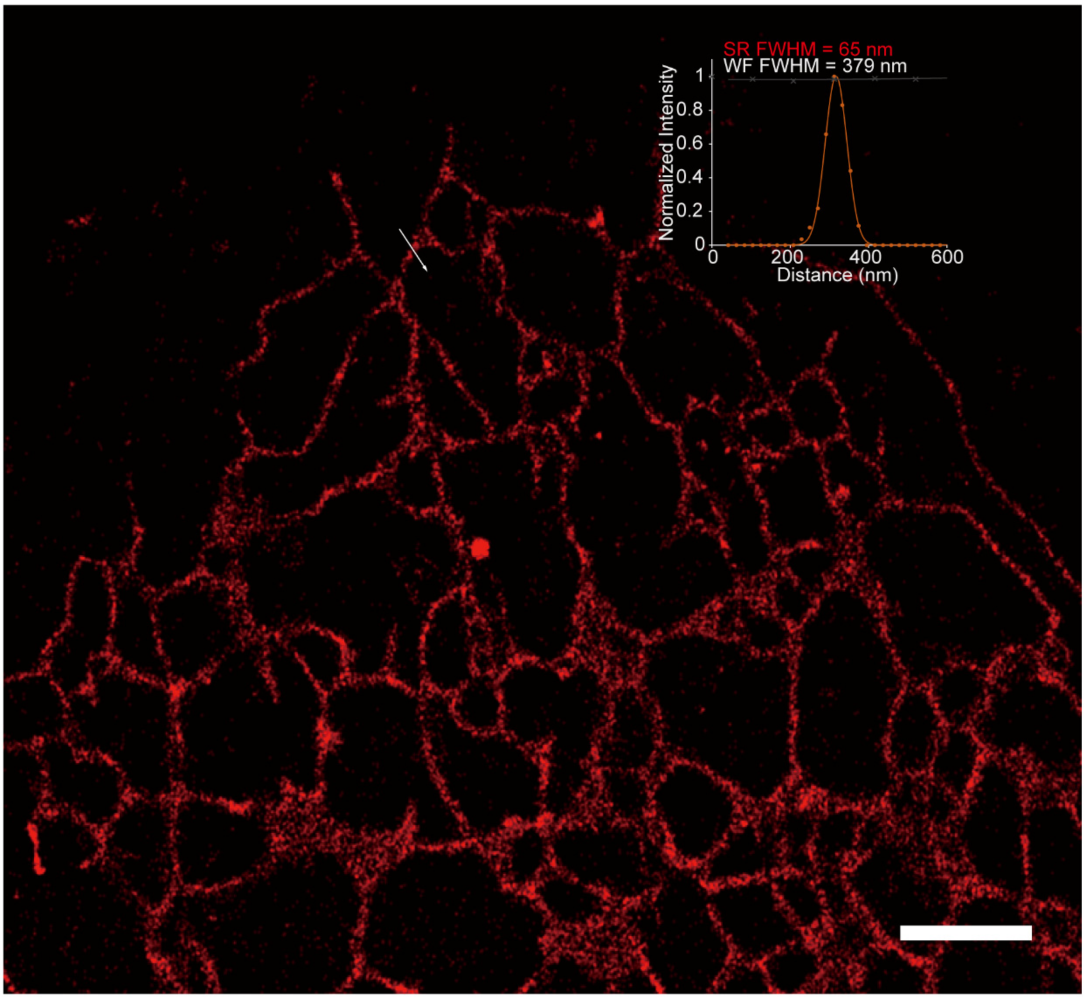

Figure S8. Super-resolution imaging of endoplasmic reticulum in living HeLa cells with SiSRh-CA. The inset represents intensity profiles of endoplasmic reticulum highlighted with white lines in the super-resolution image. Scale bars: 2 μm.

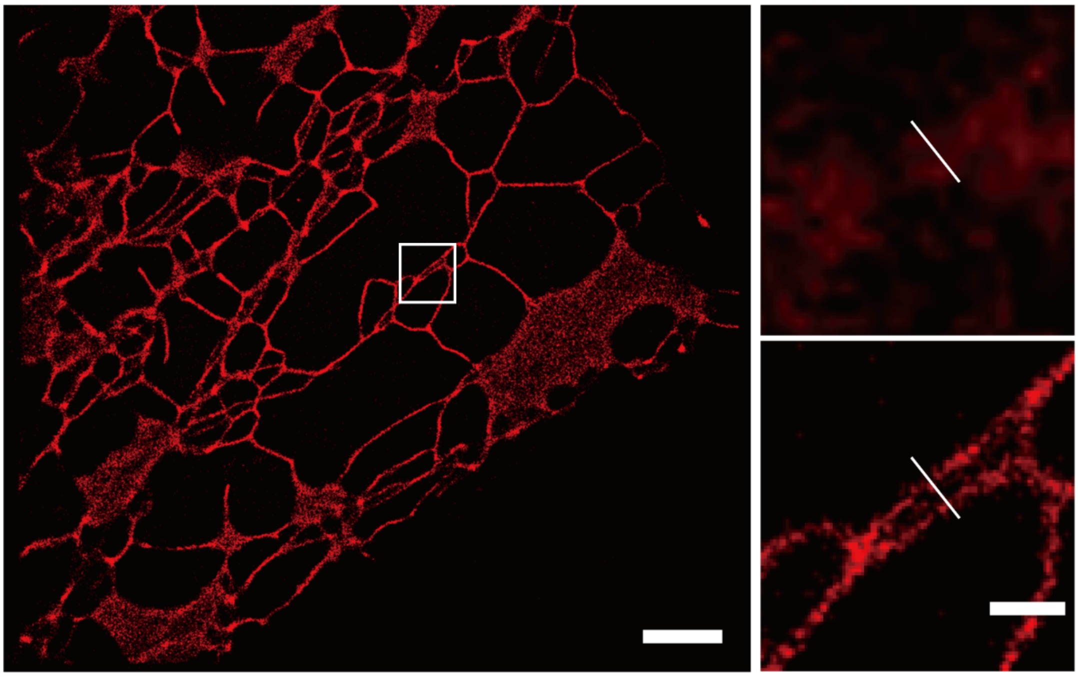

Figure S9. Super-resolution imaging of endoplasmic reticulum in living HeLa cells with CSRh-CA. Insets on the right panel shows the conventional image (WF) and the super-resolution image (SR) of a magnified view at the highlighted regions. Scale bars: 3 μm; 0.5 μm (inset).

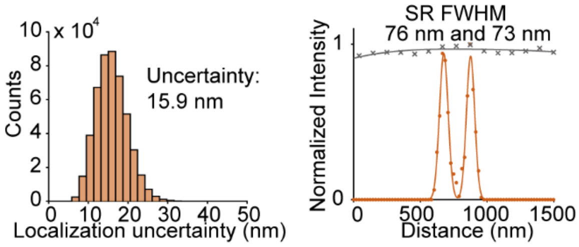

Figure S10. Histogram of localization uncertainty and intensity profiles of CSRh-CA on endoplasmic reticulum highlighted with white lines in the super-resolution image.

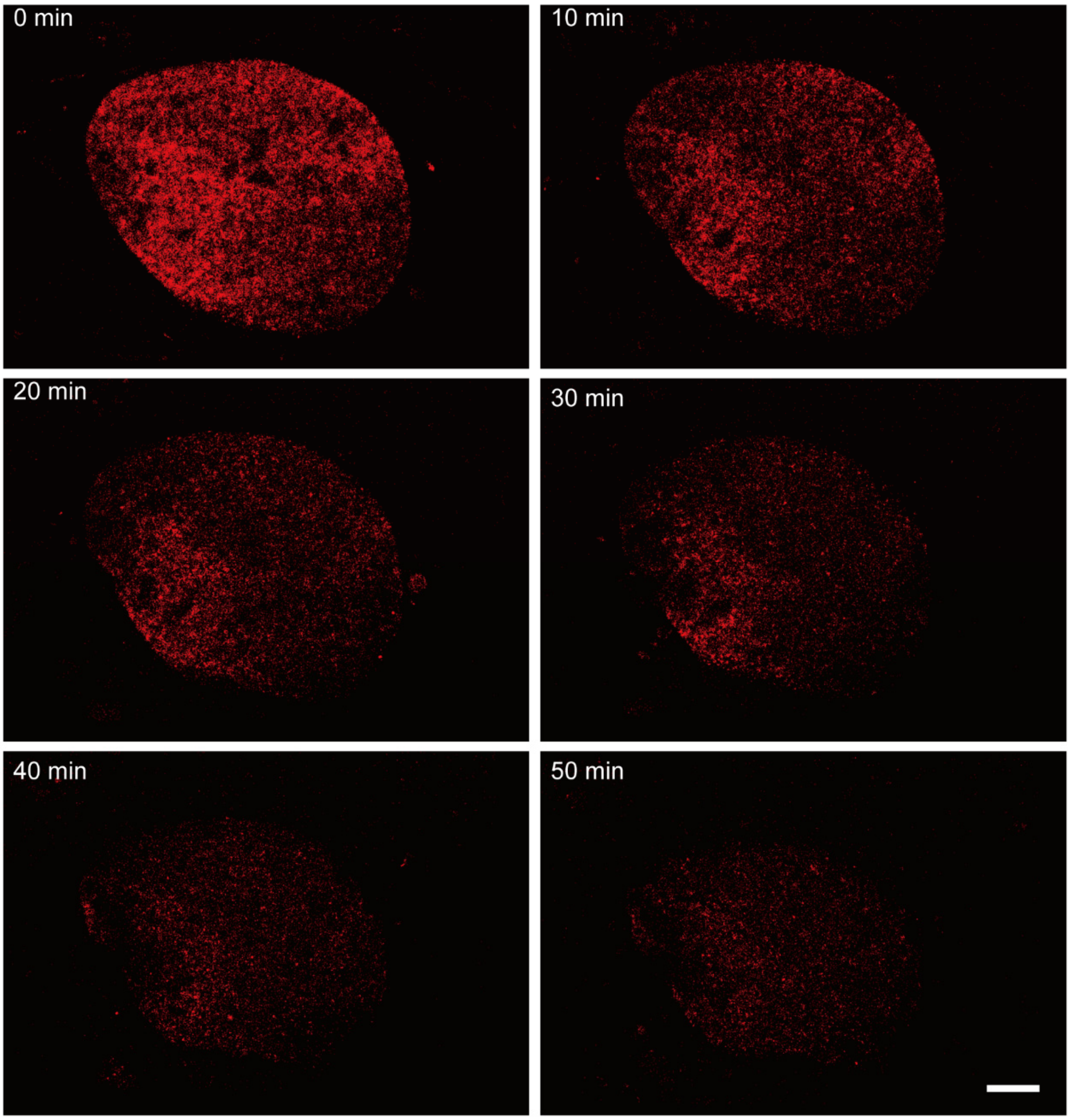

Figure S11. Long-term super-resolution imaging of H2B proteins in living U2OS cells with SiSRh-CA. Scale bar: 3 μm.

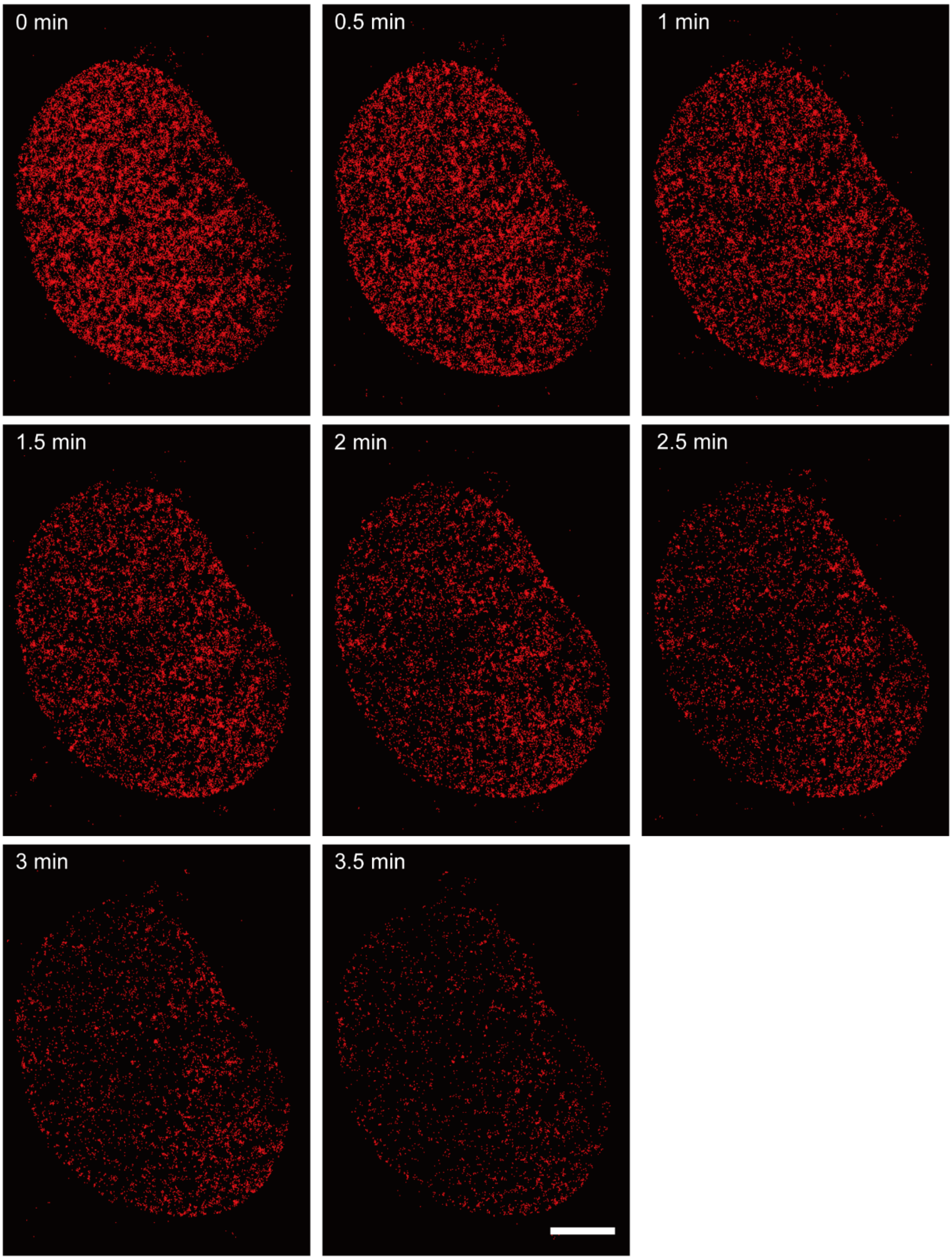

Figure S12. Long-term super-resolution imaging of H2B proteins in living U2OS cells with CSRh-CA. Scale bar: 3 μm.

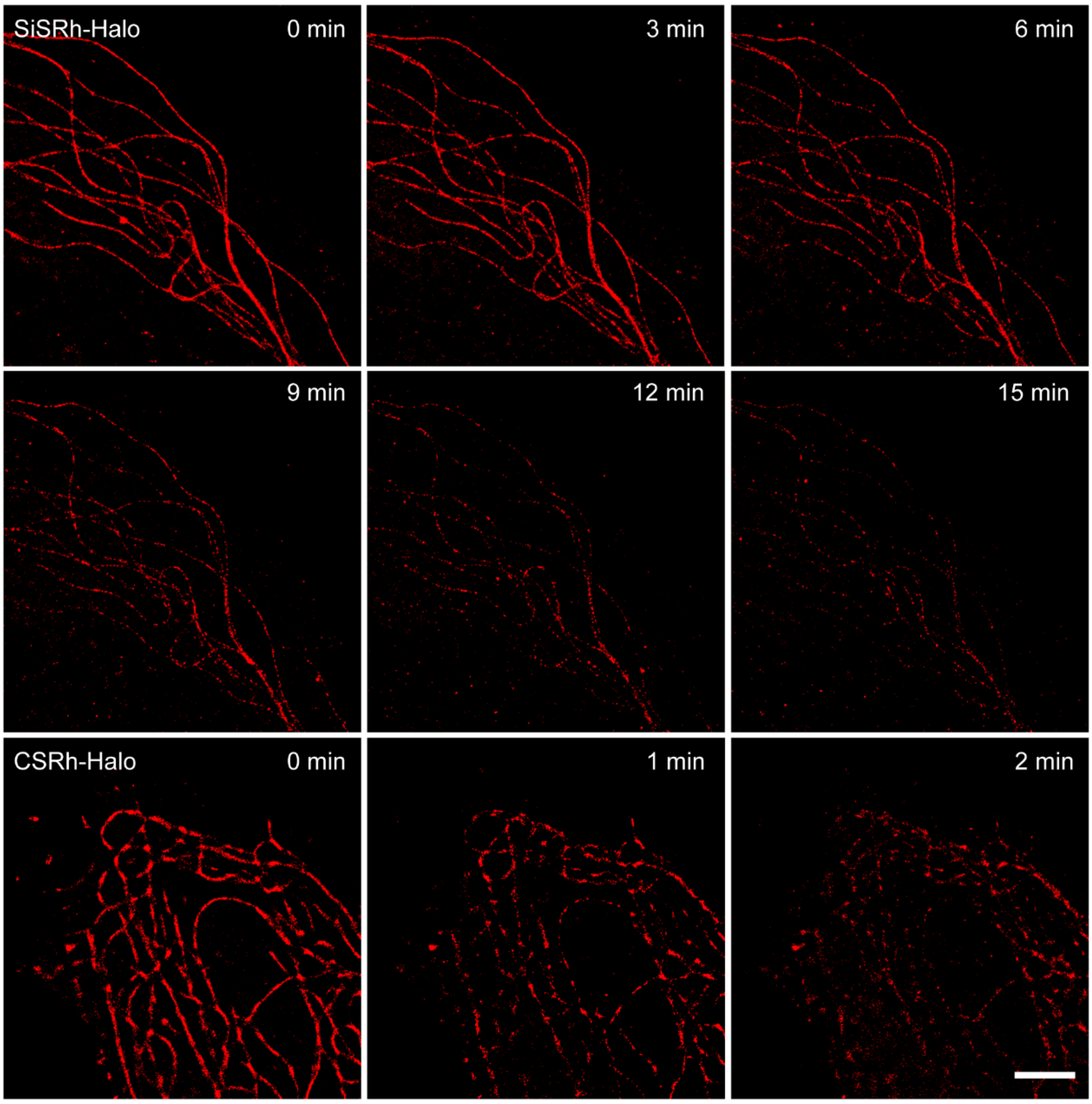

Figure S13. Long-term super-resolution imaging of microtubules in living Vero cells with SiSRh-CA and CSRh-CA. Scale bar: 3 μm.

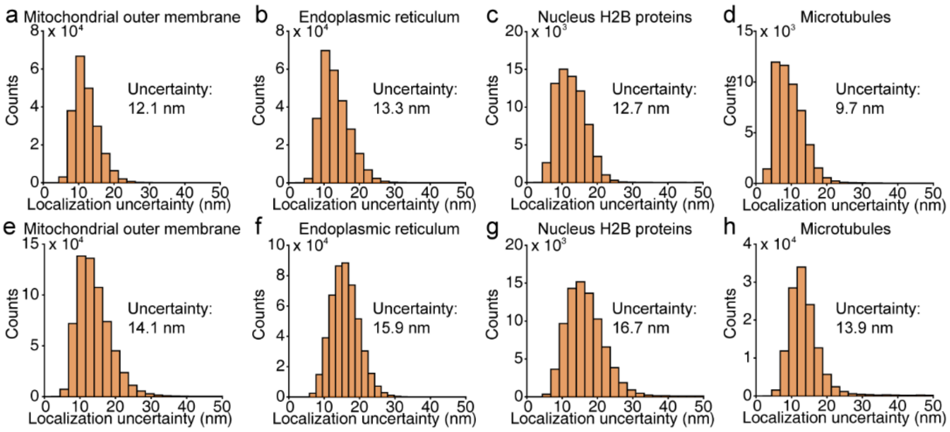

Figure S14. Histogram of localization uncertainty of SiSRh-CA (a-d) and CSRh-CA (e-h) on reconstructed cellular organelles.

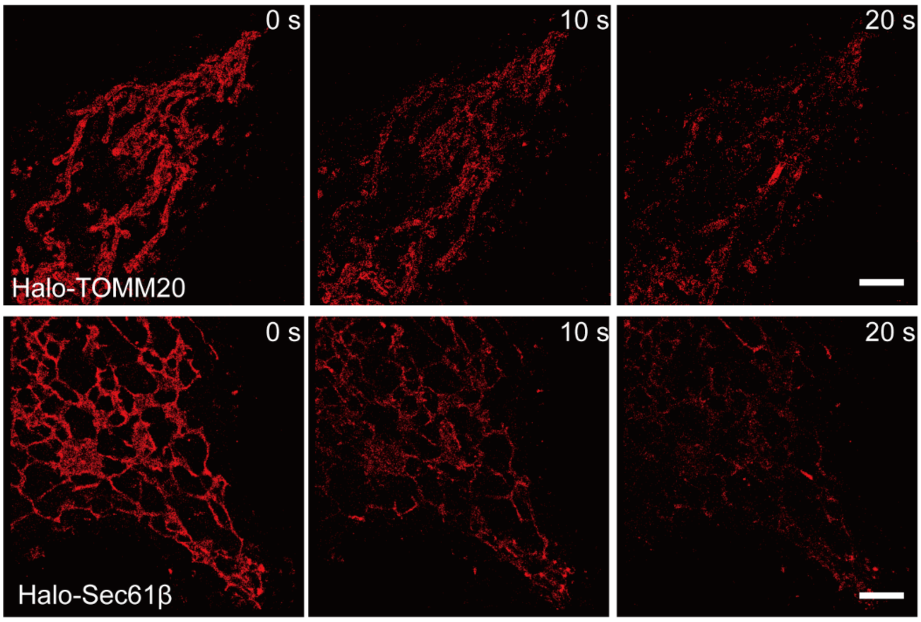

Figure S15. Long-term super-resolution imaging of mitochondrial and ER in live cell with HMSiR. Scale bars: 3 μm.

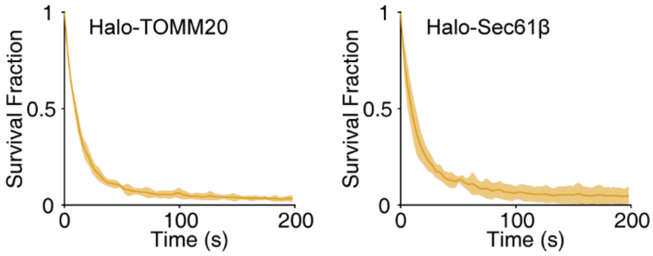

Figure S16. The time dependent survival fraction of imaging in mitochondria and endoplasmic reticulum with HMSiR.

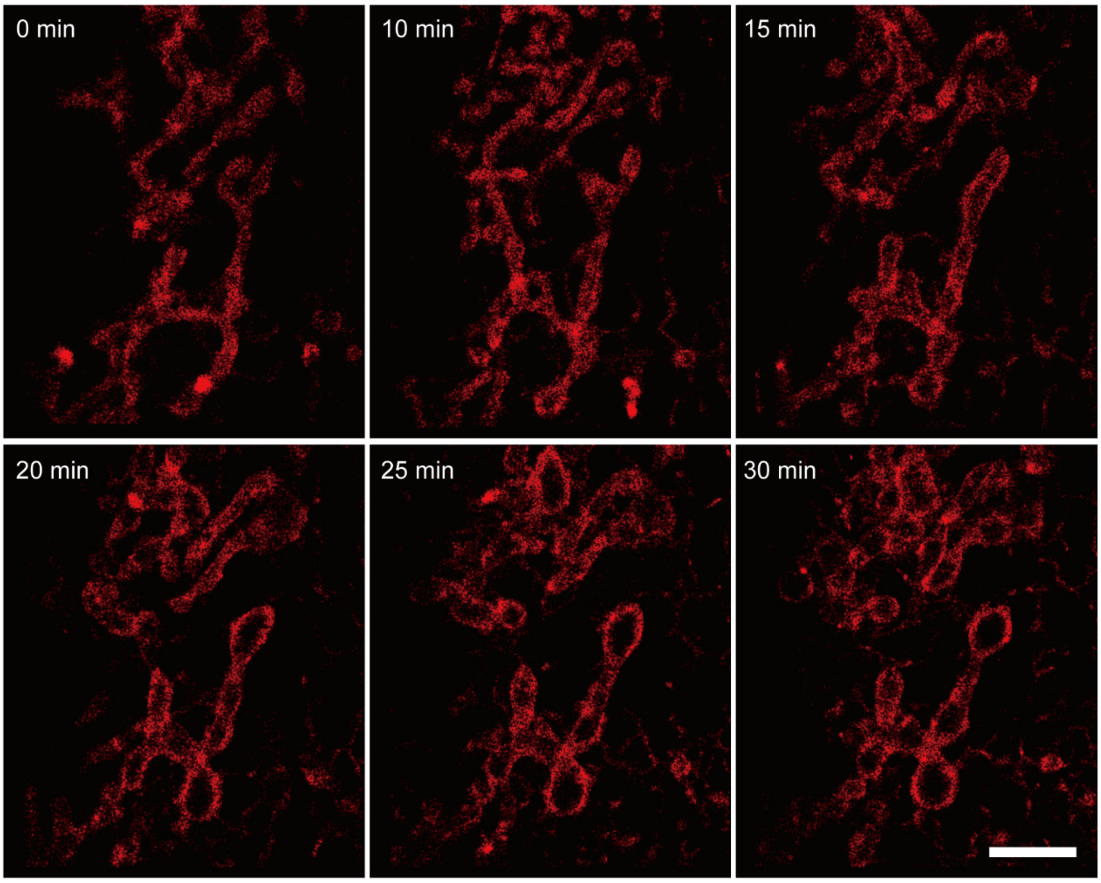

Figure S17. Long-term super-resolution imaging of mitochondrial in living U2OS cells with SiSRh-CLP. Scale bar: 3 μm.

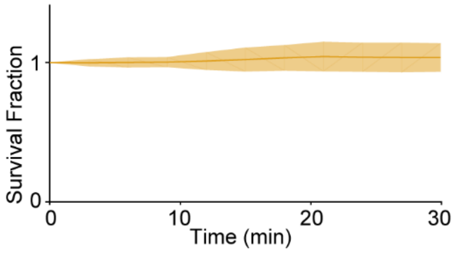

Figure S18. The time dependent survival fraction of imaging in mitochondria with SiSRh-CLP.

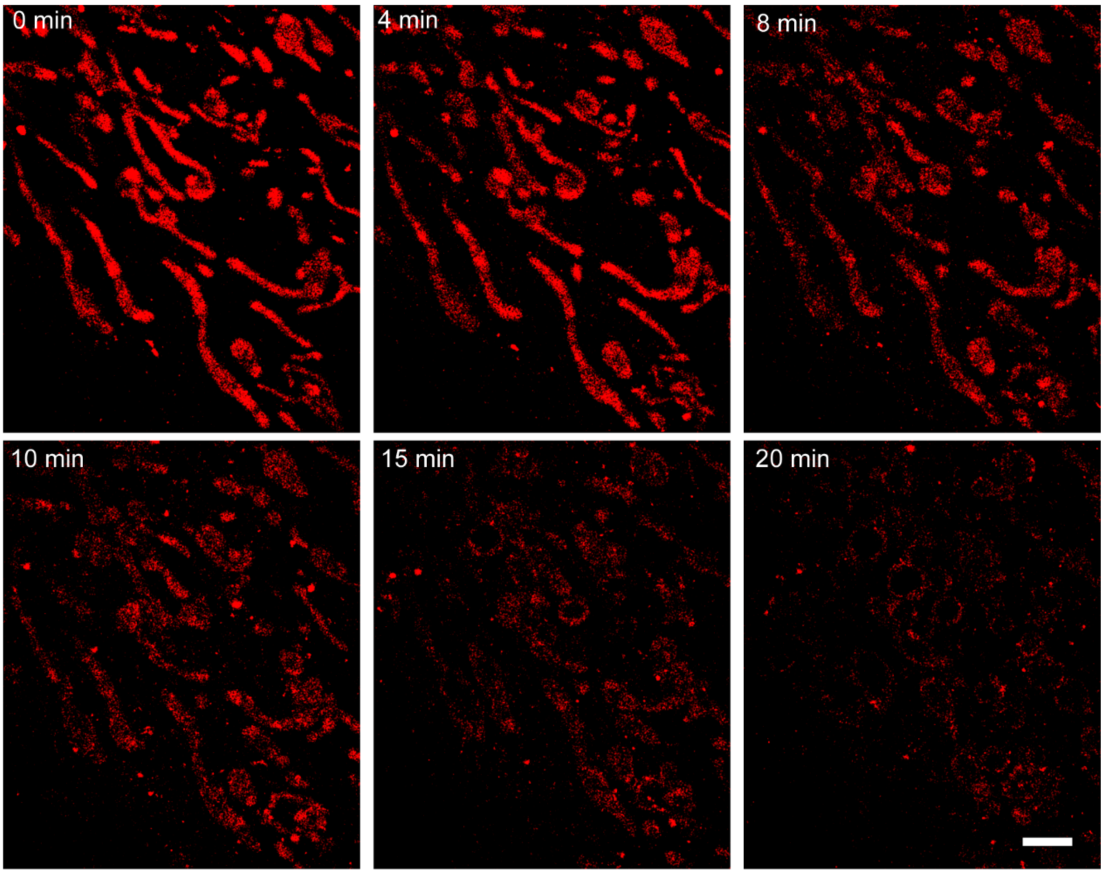

Figure S19. Long-term super-resolution imaging of mitochondrial in living Hela cells with SiSRh-Tz. Scale bar: 3 μm.

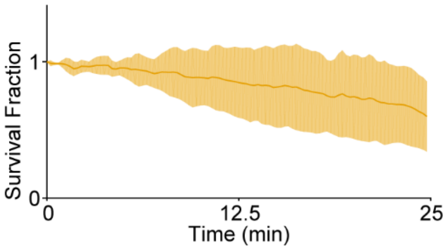

Figure S20. The time dependent survival fraction of imaging in mitochondria with SiSRh-Tz.

##### 4.4 **Microtubules in fixed cells**

For microtubule imaging in fixed cells, HeLa cells were fixed, immunostained according to a previously described procedure.[4,12]

Imaging: The imaging media was TN Buffer (50 mM Tris, pH 8.0 and 10 mM NaCl) supplemented with 200 mM cysteamine (MEA), 50 U/mL glucose oxidase, 300 U/mL catalase and 10% glucose. The super-resolution imaging was obtained with 800 W/cm$^2$ 640 nm laser. The imaging of SiSRh and AF647 was acquired in 2000 frames.

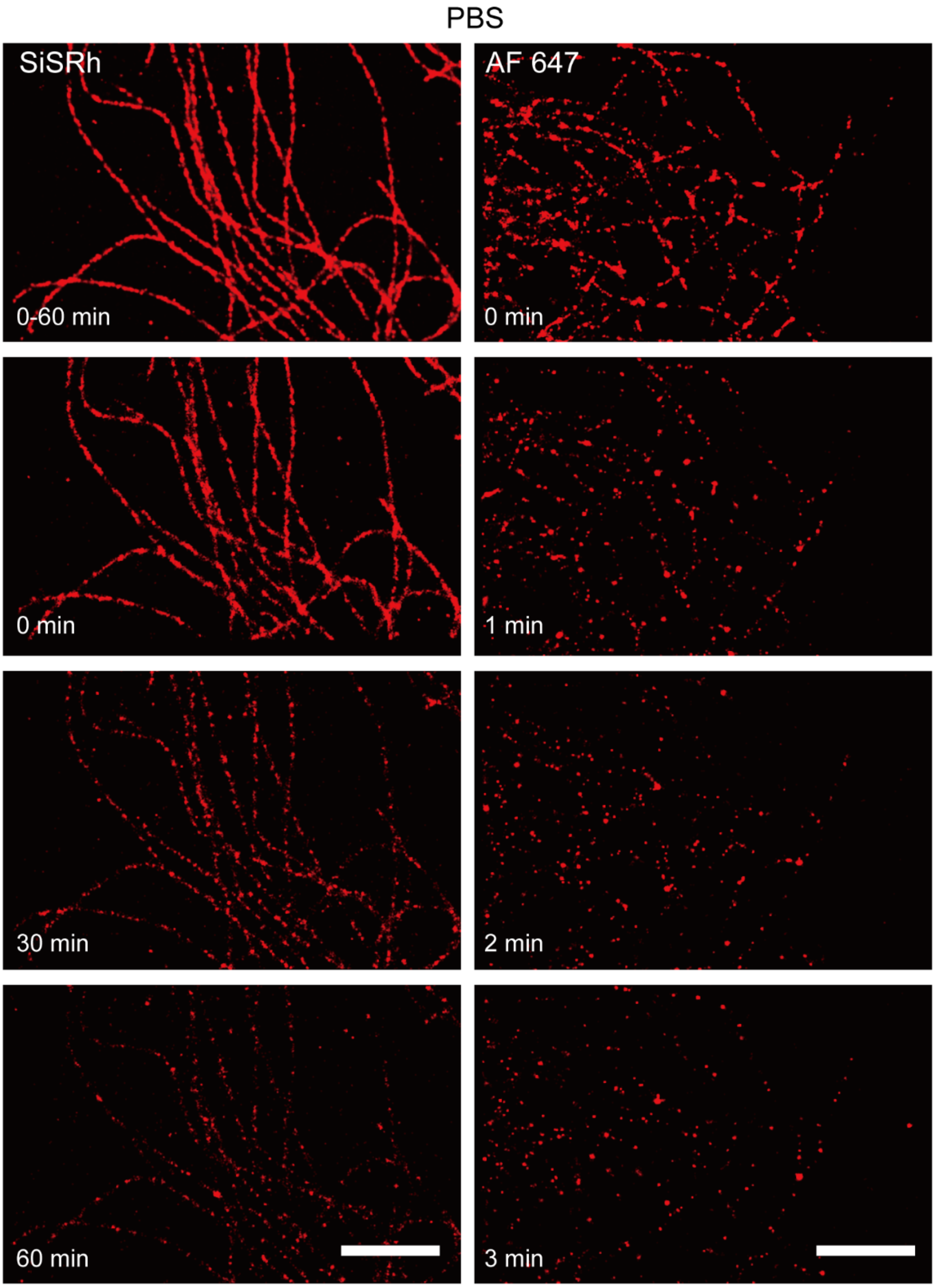

Figure S21. The time-lapse images of fixed cell microtubules in PBS with SiSRh and AF647. Scale bars: 3 μm.

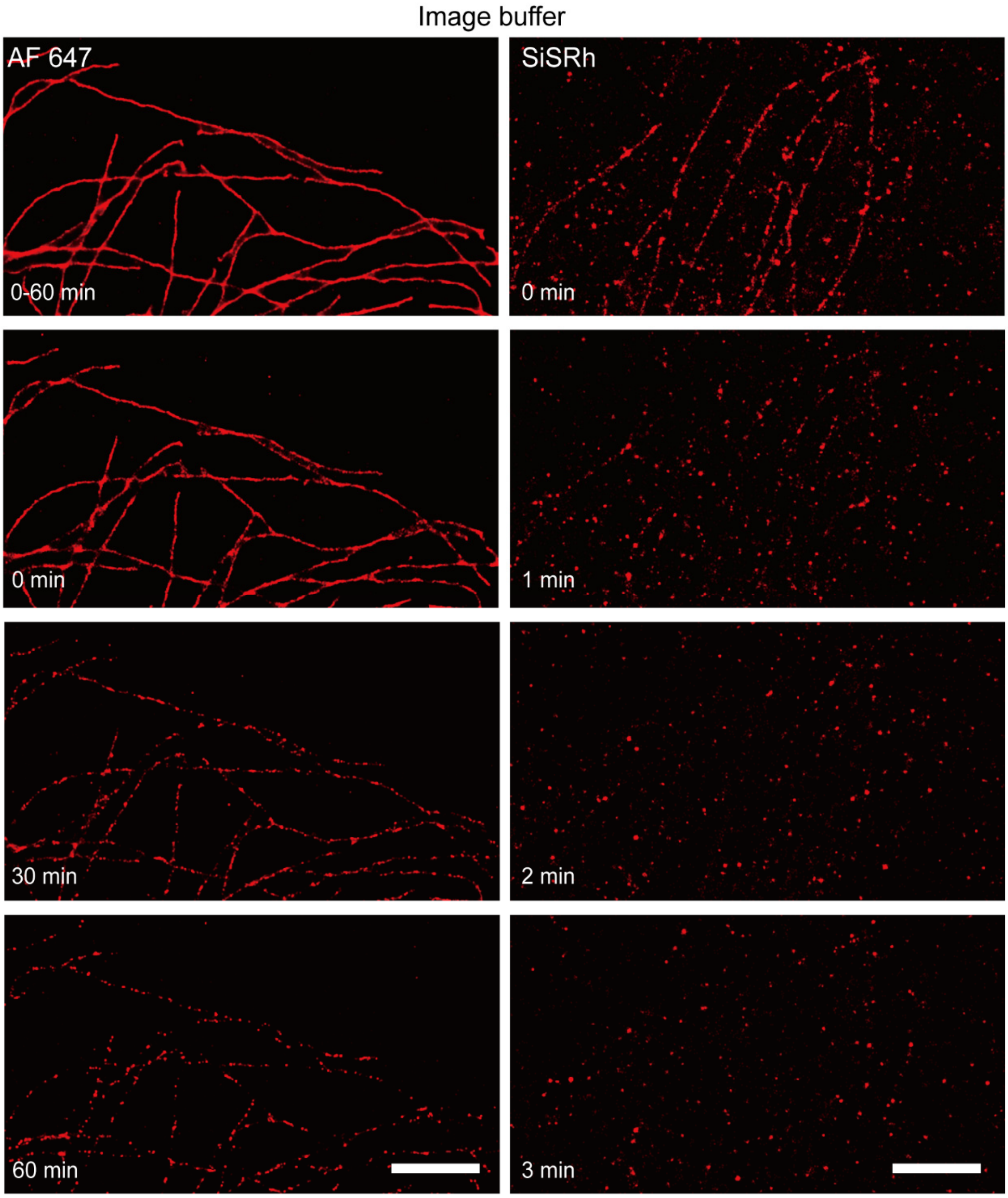

Figure S22. The time-lapse images of fixed cell microtubules in image buffer with SiSRh and AF647. Scale bars: 3 μm.

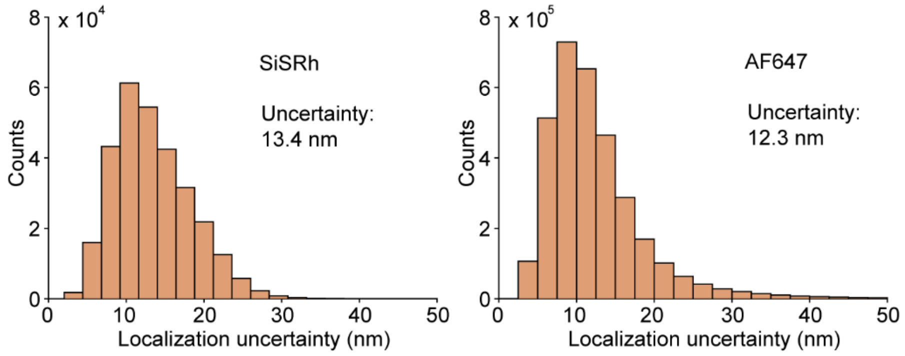

Figure S23. Histogram of localization uncertainty of SiSRh in PBS and AF647 in image buffer.

4.5 **Long-term Imaging under stressed living cell condition**

Super-resolution imaging acquisition under stressed living cell condition. Halo-TOMM20-expressing cells were incubated with 200 nM SiSRh-CA were stained in cells for 2 h. The free remaining dyes were washed with PBS for three times and were further cultured in a $CO_2$ incubator with fresh McCoy's 5A(PM150710) media for 90 min. DUT870 (600 nM) was stained for another 20 min and irradiated with 808 nm laser (0.33 W/cm$^2$) for 5 min. The imaging media was McCoy's 5A(PM150710) without phenol red supplemented with 10% FBS.

4.6 **Dual-color SMLM image of live U2OS cells**

Halo-TOMM20 and SNAP-ER expressing cells were incubated with 500 nM SiSRh-CA, 200 nM STMR-CLP. Halo-TOMM20 and SNAP-COX8A expressing cells were incubated with 300 nM SiSRh-CA, and 100 nM STMR-CLP.

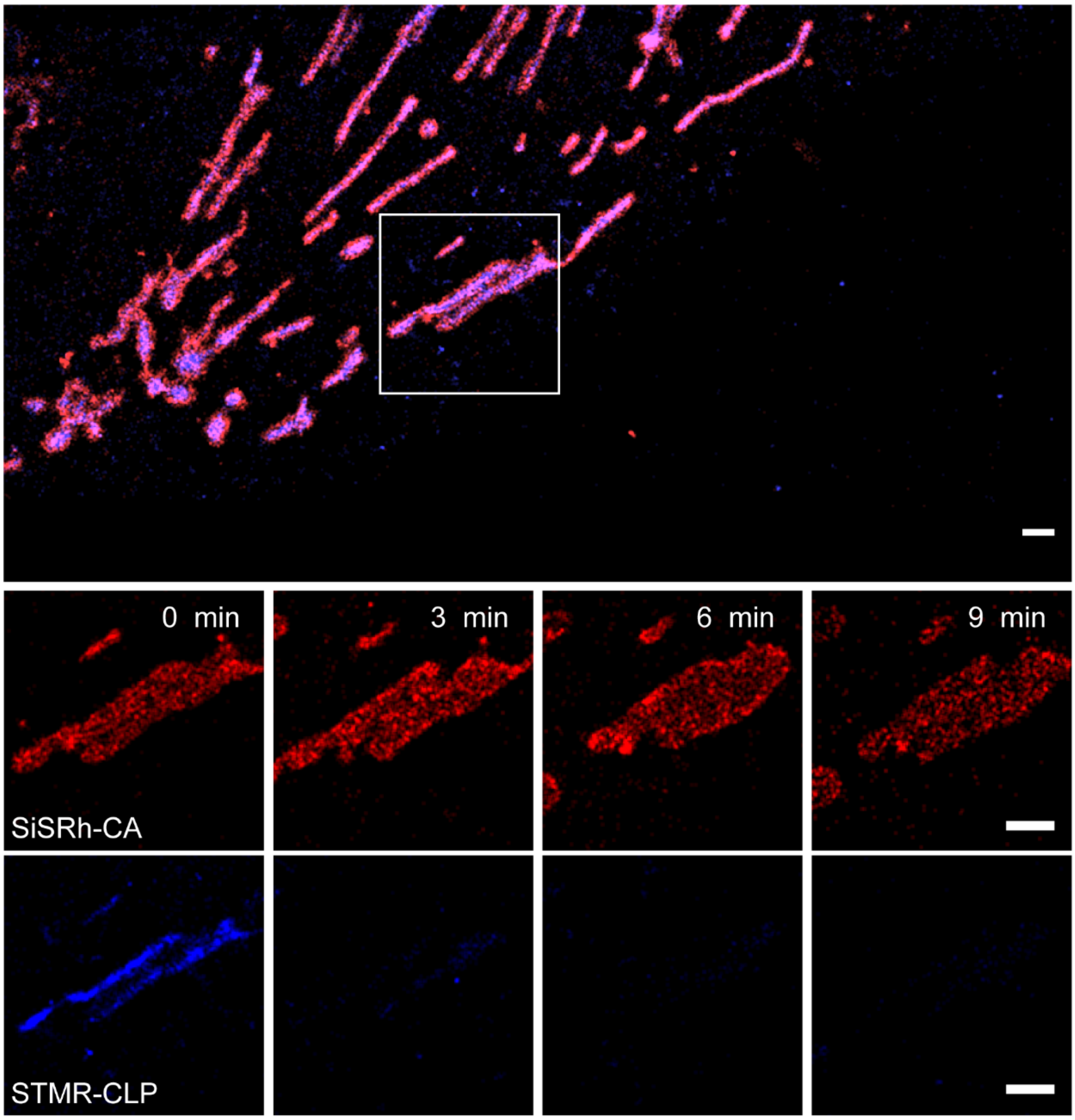

Figure S24. Long-term dual-color SMLM image of U2OS cells expressing Halo-TOMM20 (mitochondrial outer membrane, treated with SiSRh) and SNAP-COX8A (mitochondrial inner membrane, treated with STMR). Scale bars: 1 μm.

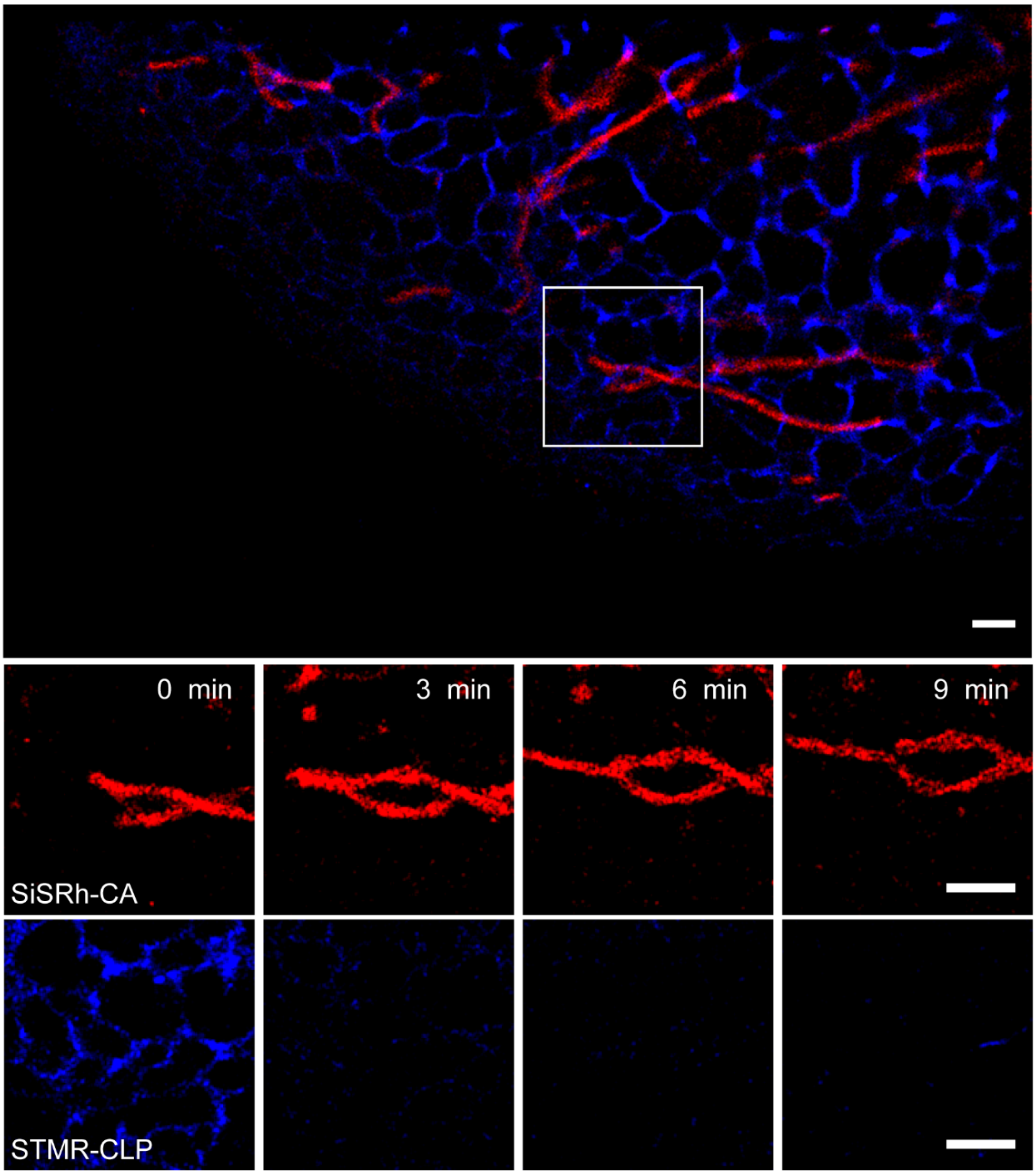

Figure S25. Long-term dual-color single-molecule super-resolution image of U2OS cells expressing Halo-TOMM20 (mitochondrial outer membrane, treated with SiSRh) and SNAP-ER (endoplasmic reticulum, treated with STMR). Scale bars: 1 μm.

#### 5 Movies descriptions

Movie S1. The 30 min movie of the mitochondrial in U2OS cells treated with SiSRh-CA was performed with 800 W/cm$^2$ 640 nm laser. The exposure time for single image was 10 ms.

Movie S2. The 30 min movie of the ER in Hela cells treated with SiSRh-CA was performed with 800 W/cm$^2$ 640 nm laser. The exposure time for single image was 10 ms.

Movie S3. The 10 min movie of the mitochondrial in U2OS cells treated with CSRh-CA was performed with 800 W/cm$^2$ 640 nm laser. The exposure time for single image was 10 ms.

Movie S4. The 10 min movie of the ER in Hela cells treated with CSRh-CA was performed with 800 W/cm$^2$ 640 nm laser. The exposure time for single image was 10 ms.

Movie S5. The 60 min movie of the microtubules in fixed Hela cells treated with SiSRh-COOH was performed with 800 W/cm$^2$ 640 nm laser. The exposure time for single image was 50 ms.

#### 6 Characterization spectra

##### 6.1 $^1$H NMR spectrum of 1

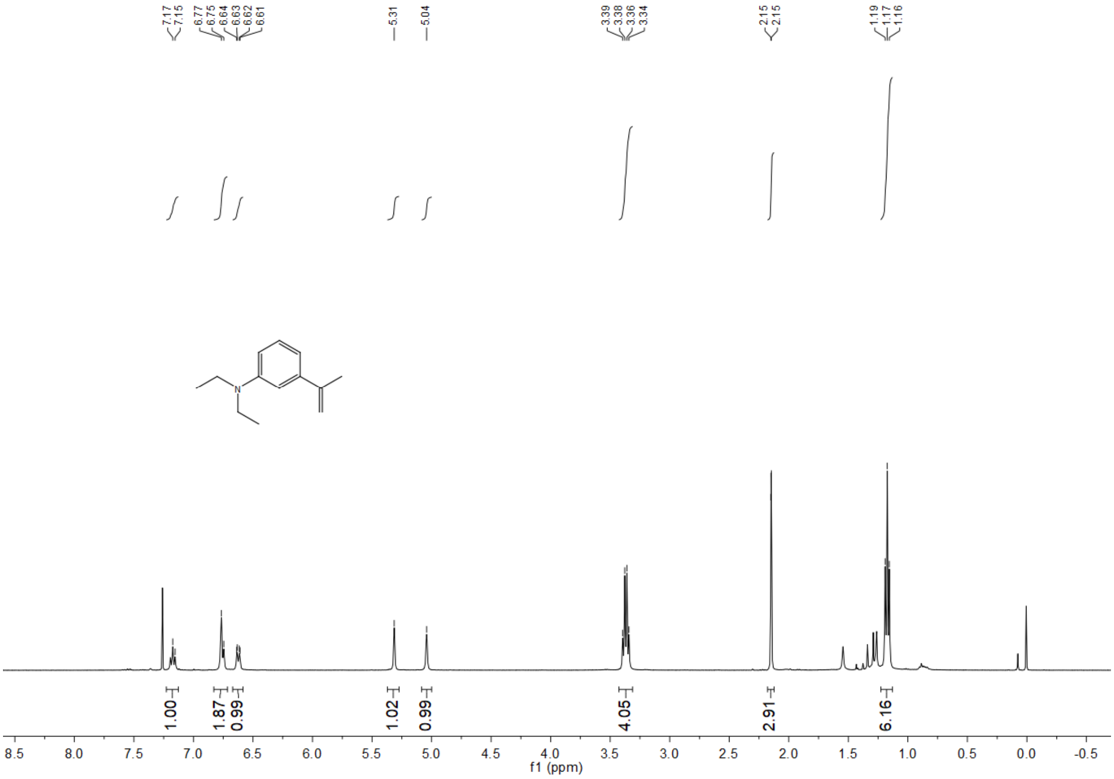

#### 6.2 $^{1}$H NMR spectrum of 3

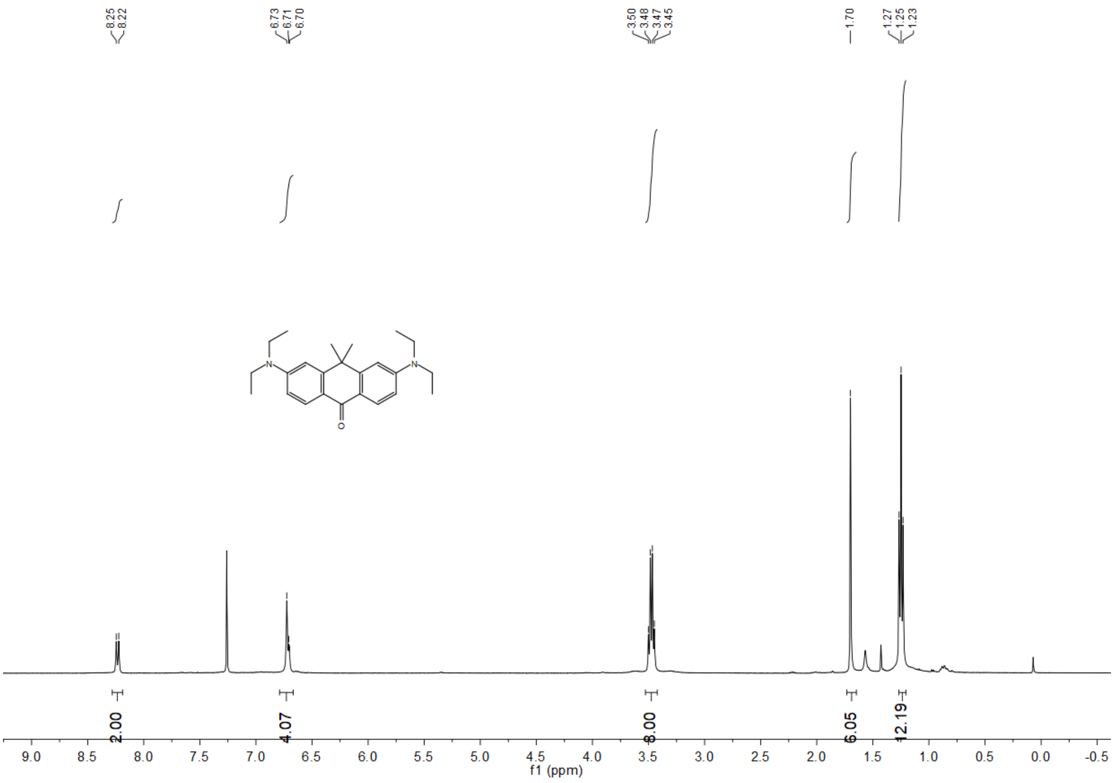

#### 6.3 $^{1}$H NMR spectrum of CSRh

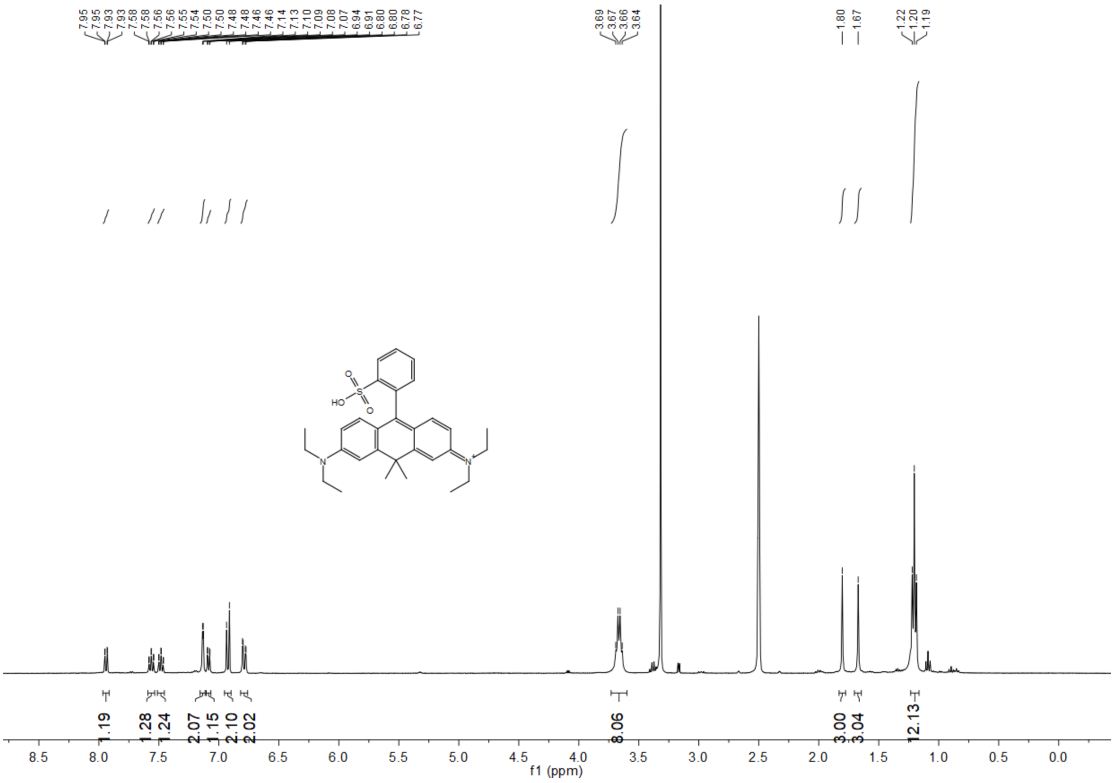

#### 6.4 $^{13}$C NMR spectrum of CSRh

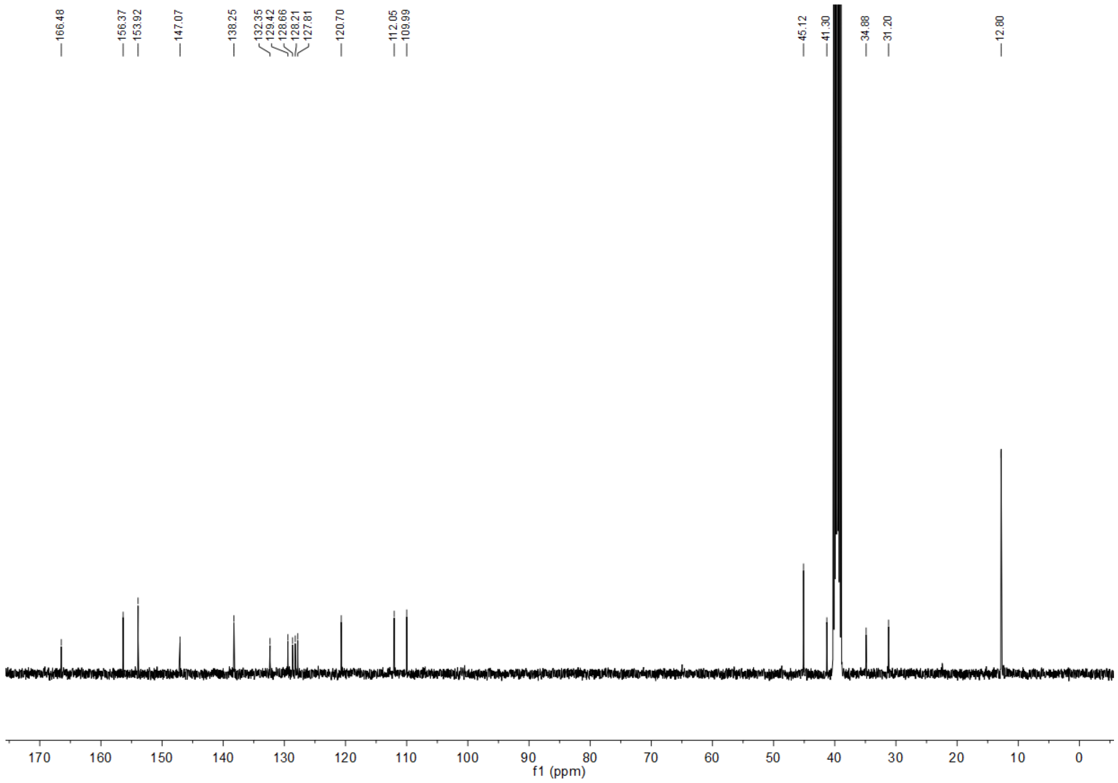

#### 6.5 $^{1}$H NMR spectrum of CSRh-COOH

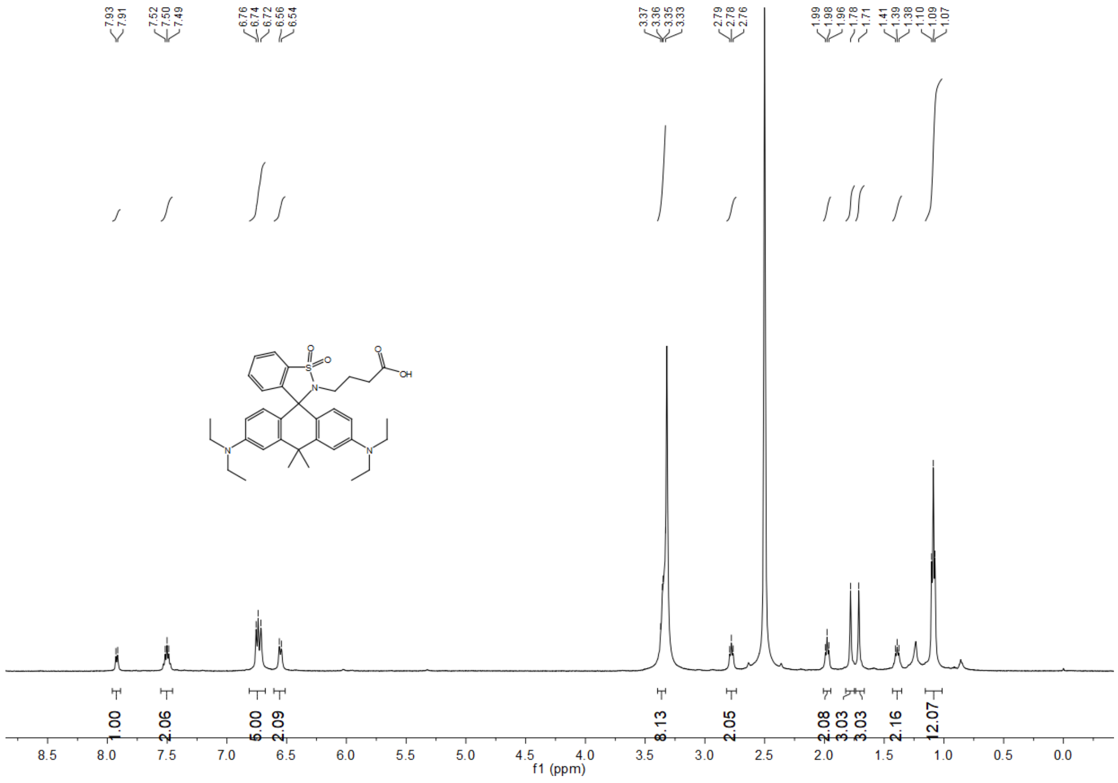

#### 6.6 $^{13}$C NMR spectrum of CSRh-COOH

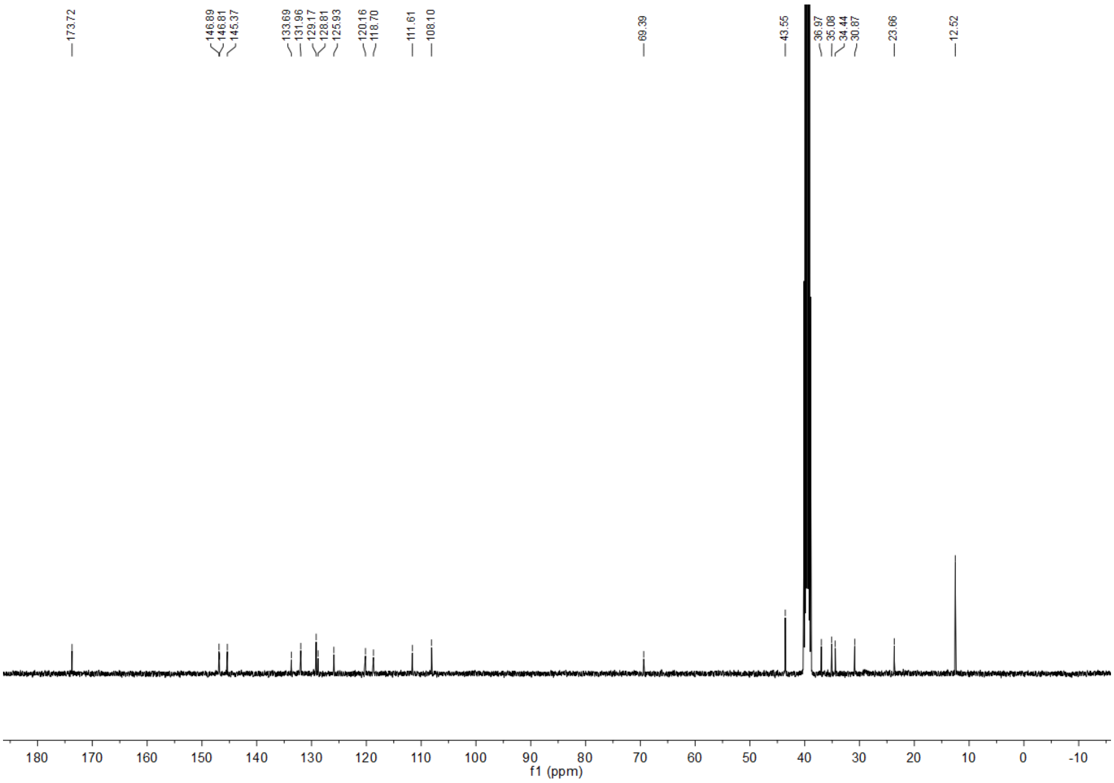

#### 6.7 $^{1}$H NMR spectrum of CSRh-CA

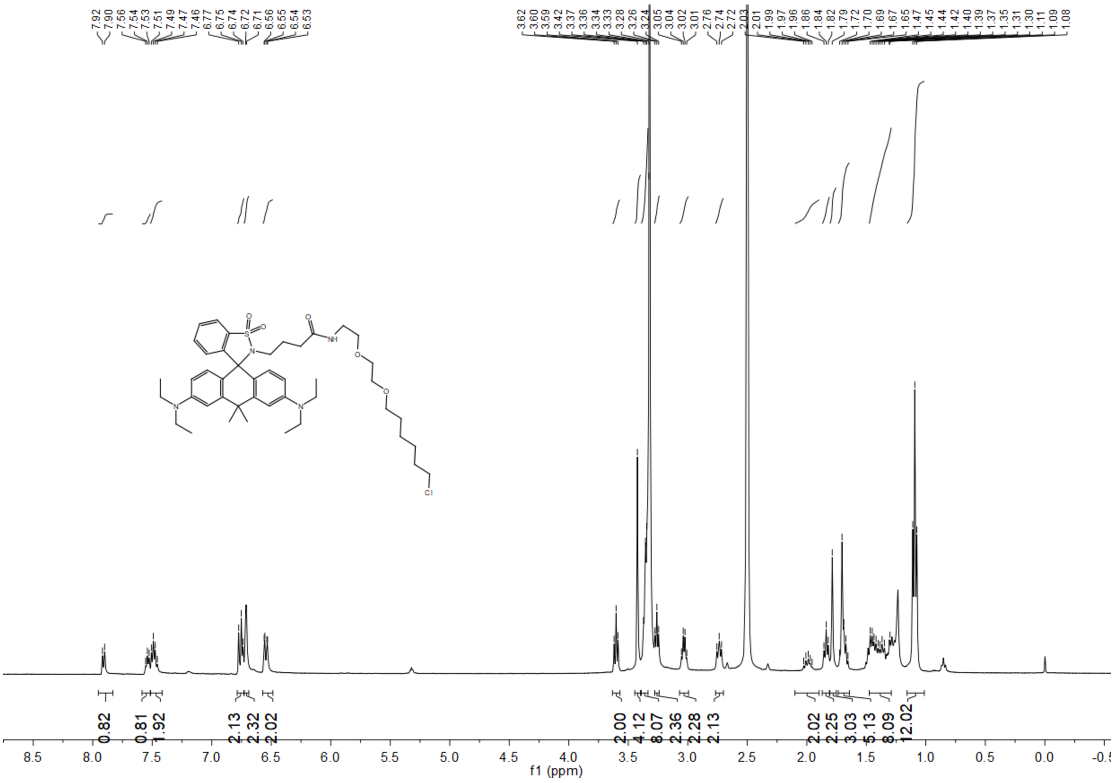

#### 6.8 $^{13}C$ NMR spectrum of CSRh-CA

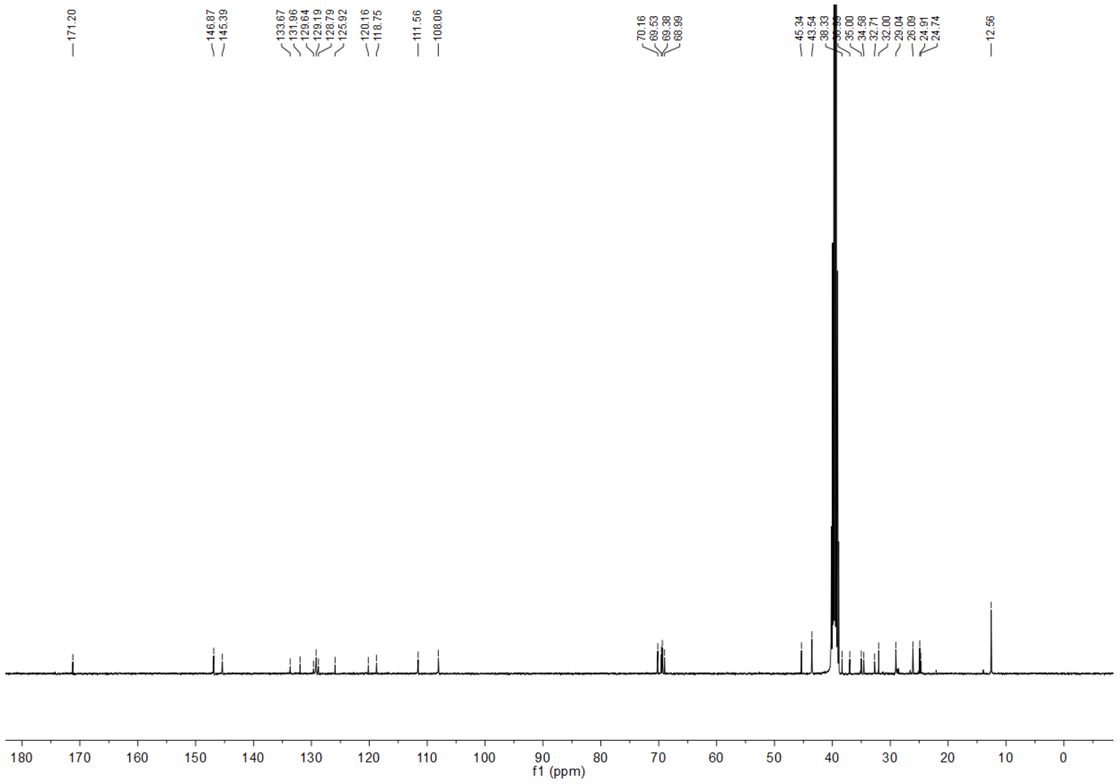

#### 6.9 $^{1}H$ NMR spectrum of SiSRh

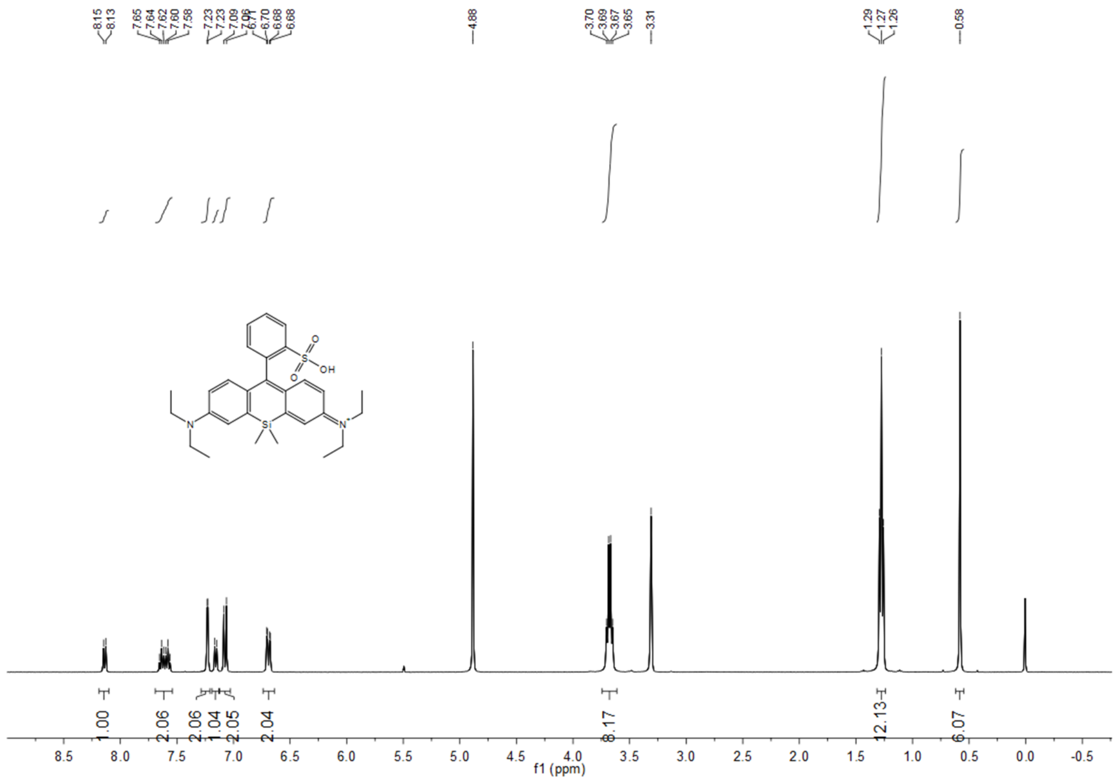

#### 6.10 $^{13}C$ NMR spectrum of SiSRh

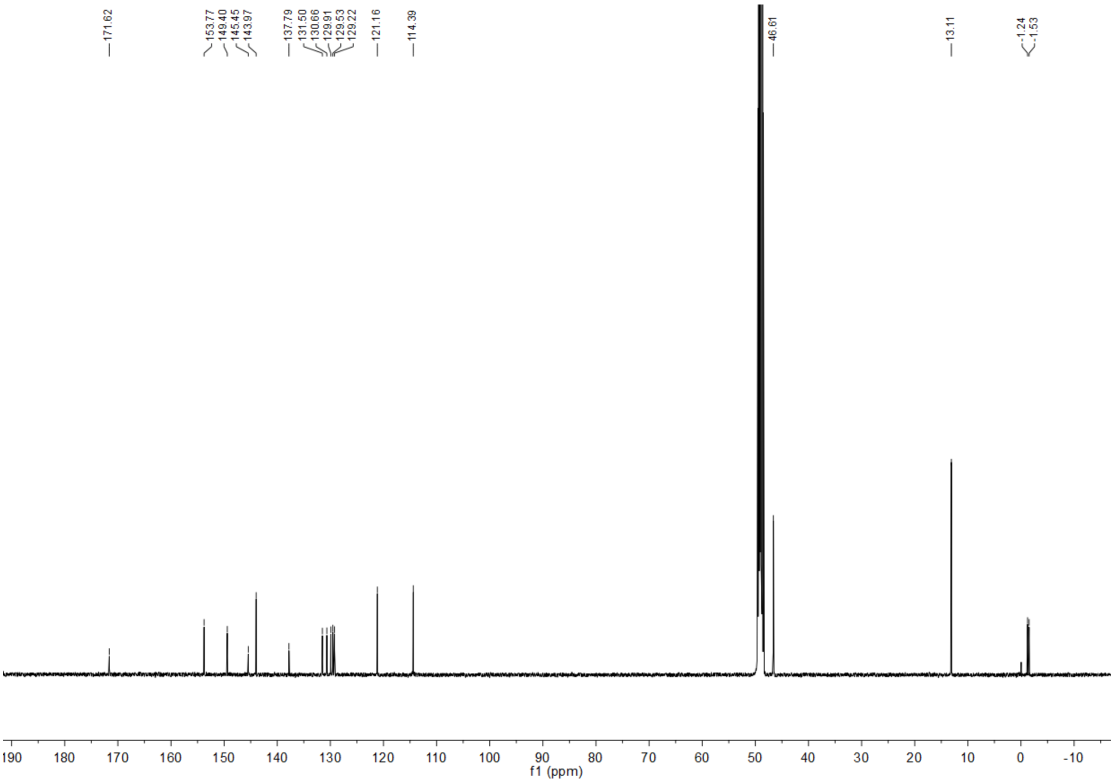

#### 6.2 $^{1}H$ NMR spectrum of SiSRh-COOH

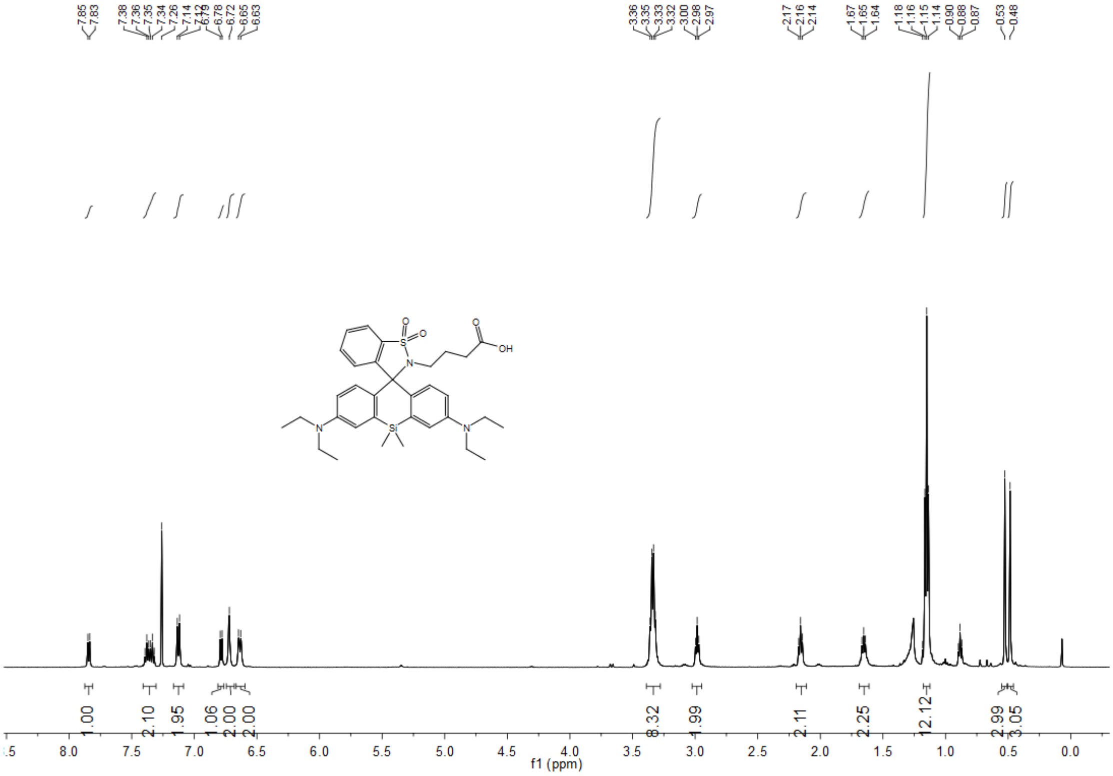

#### 6.3 $^{1}H$ NMR spectrum of SiSRh-CA

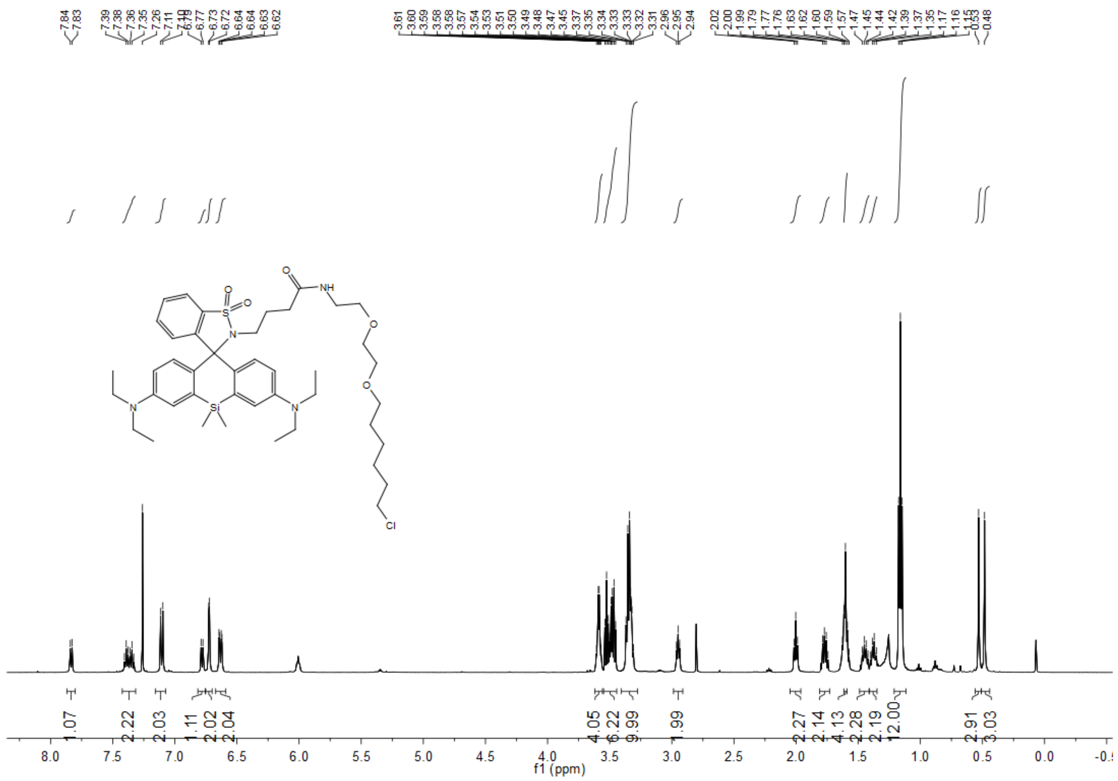

#### 6.4 $^{13}C$ NMR spectrum of SiSRh-CA

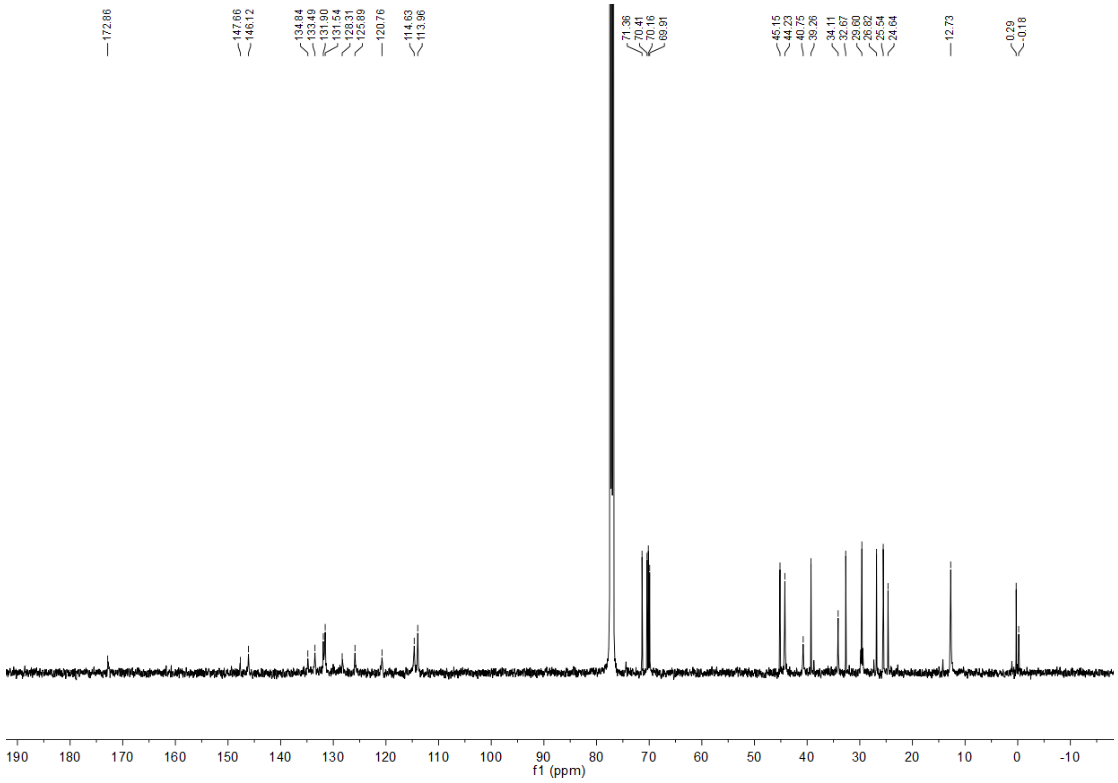

#### 6.5 $^{1}$H NMR spectrum of SiSRh-CLP

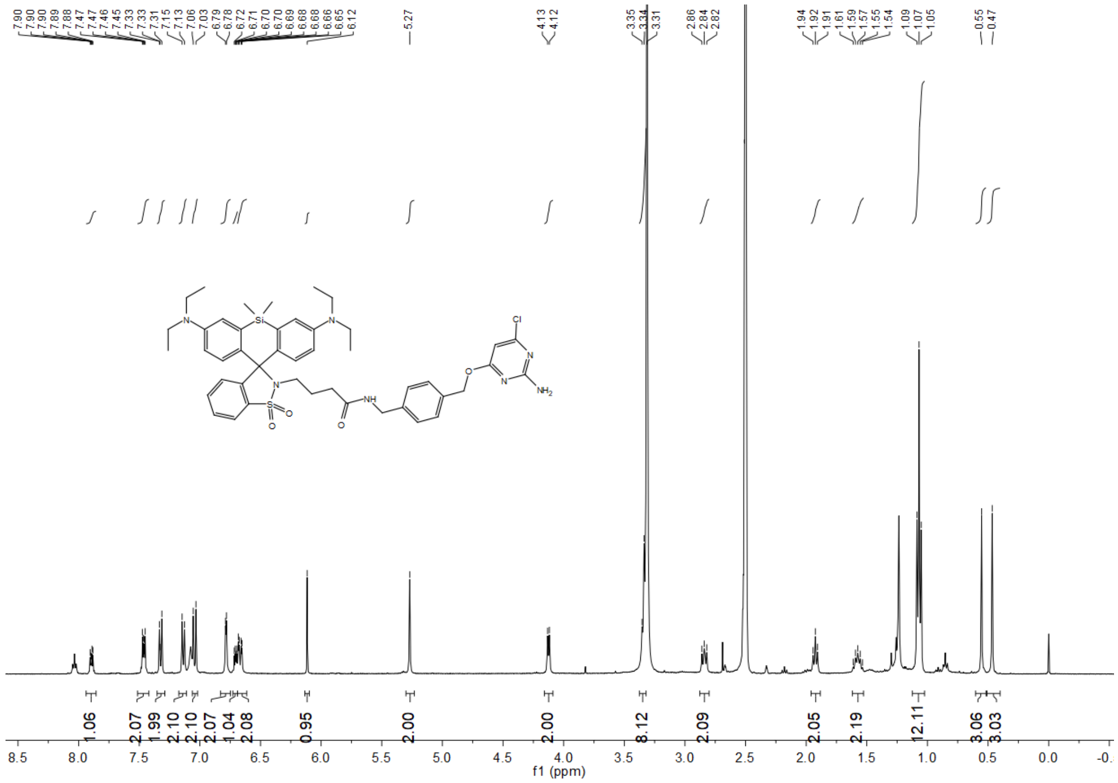

#### 6.6 $^{13}$C NMR spectrum of SiSRh-CLP

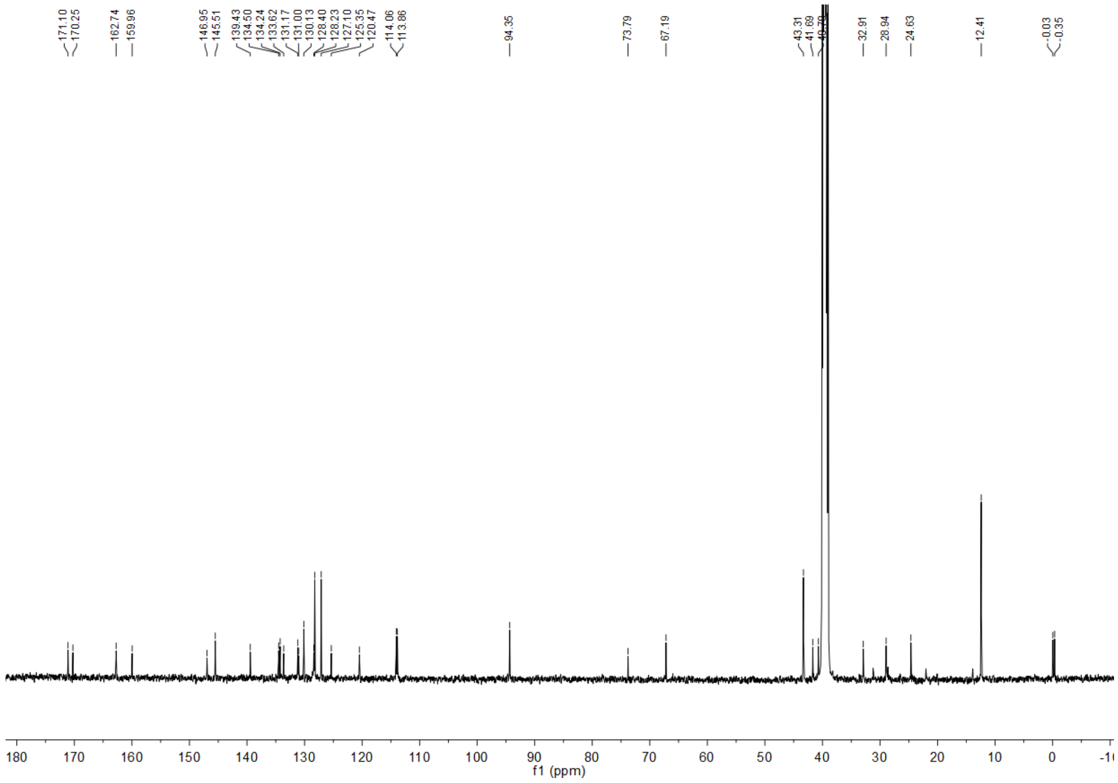

#### 6.7 $^{1}$H NMR spectrum of SiSRh-Tz

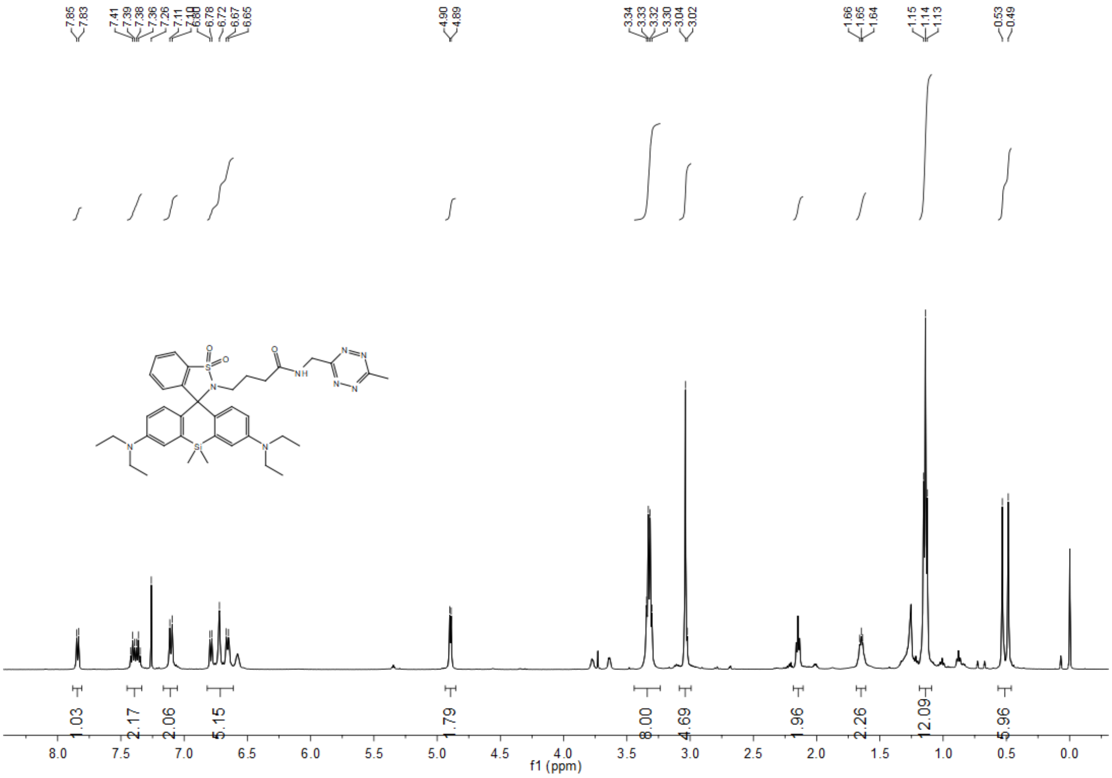

### 7 Reference

1. Bucevičius, J.; Gerasimaitė, R.; Kiszka, K. A.; Pradhan, S.; Kostiuk, G.; Koenen, T.; Lukinavičius, G. A General Highly Efficient Synthesis of Biocompatible Rhodamine Dyes and Probes for Live-Cell Multicolor Nanoscopy. Nat. Commun. 2023, 14 (1), 1–14.

2. Grimm, J. B.; Tkachuk, A. N.; Patel, R.; Hennigan, S. T.; Gutu, A.; Dong, P.; Gandin, V.; Osowski, A. M.; Holland, K. L.; Liu, Z. J.; Brown, T. A.; Lavis, L. D. Optimized Red-Absorbing Dyes for Imaging and Sensing. J. Am. Chem. Soc. 2023.

3. Ye, Z.; Yang, W.; Wang, C.; Zheng, Y.; Chi, W.; Liu, X.; Huang, Z.; Li, X.; Xiao, Y., Quaternary Piperazine-Substituted Rhodamines with Enhanced Brightness for Super-Resolution Imaging. J. Am. Chem. Soc. 2019, 141 (37), 14491-14495.

4. Ye, Z.; Yu, H.; Yang, W.; Zheng, Y.; Li, N.; Bian, H.; Wang, Z.; Liu, Q.; Song, Y.; Zhang, M.; Xiao, Y., Strategy to Lengthen the On-Time of Photochromic Rhodamine Spirolactam for Super- resolution Photoactivated Localization Microscopy. J. Am. Chem. Soc. 2019, 141 (16), 6527-6536.

5. He, H.; Ye, Z.; Xiao, Y.; Yang, W.; Qian, X.; Yang, Y., Super-Resolution Monitoring of Mitochondrial Dynamics upon Time-Gated Photo-Triggered Release of Nitric Oxide. Anal. Chem. 2018, 90 (3), 2164-2169.

6. Ovesný, M.; Křížek, P.; Borkovec, J.; Švindrych, Z.; Hagen, G. M., ThunderSTORM: a comprehensive ImageJ plug-in for PALM and STORM data analysis and super-resolution imaging. Bioinformatics 2014, 30 (16), 2389-2390.

7. Schneider, C. A.; Rasband, W. S.; Eliceiri, K. W., NIH Image to ImageJ: 25 years of image analysis. Nat. Methods 2012, 9 (7), 671-675.

8. Mortensen, K. I.; Churchman, L. S.; Spudich, J. A.; Flyvbjerg, H., Optimized localization analysis for single-molecule tracking and super-resolution microscopy. Nat. Methods 2010, 7 (5), 377- 381.

9. Deschout, H.; Zanacchi, F. C.; Mlodzianoski, M.; Diaspro, A.; Bewersdorf, J.; Hess, S. T.; Braeckmans, K., Precisely and accurately localizing single emitters in fluorescence microscopy. Nat. Methods 2014, 11 (3), 253-266.

10. Thompson, R. E.; Larson, D. R.; Webb, W. W., Precise Nanometer Localization Analysis for Individual Fluorescent Probes. Biophys. J. 2002, 82 (5), 2775-2783.

11. Man, H.; Bian, H.; Zhang, X.; Wang, C.; Huang, Z.; Yan, Y.; Ye, Z.; Xiao, Y. Hybrid Labeling System for DSTORM Imaging of Endoplasmic Reticulum for Uncovering Ultrastructural Transformations under Stress Conditions. Biosens. Bioelectron. 2021, 189, 113378.

12. Ouyang, W.; Aristov, A.; Lelek, M.; Hao, X.; Zimmer, C. Deep Learning Massively

Accelerates Super-Resolution Localization Microscopy. Nat. Biotechnol. 2018, 36, 460.
